# SARS-CoV-2 Variant Spike and accessory gene mutations alter pathogenesis

**DOI:** 10.1101/2022.05.31.494211

**Authors:** M.E. McGrath, Y. Xue, C. Dillen, L. Oldfield, N. Assad-Garcia, J. Zaveri, N. Singh, L. Baracco, L. Taylor, S. Vashee, M. Frieman

**Affiliations:** Department of Microbiology and Immunology, University of Maryland School of Medicine, Baltimore, MD, 21201 USA; J. Craig Venter Institute, La Jolla, CA, 92037 and Rockville, MD, 20850 USA; Center for Pathogen Research, University of Maryland School of Medicine, Baltimore, MD, 21201 USA

**Keywords:** Coronavirus, Pathogenesis, Variant, SARS-CoV-2

## Abstract

The ongoing COVID-19 pandemic is a major public health crisis. Despite the development and deployment of vaccines against SARS-CoV-2, the pandemic persists. The continued spread of the virus is largely driven by the emergence of viral variants, which can evade the current vaccines through mutations in the Spike protein. Although these differences in Spike are important in terms of transmission and vaccine responses, these variants possess mutations in the other parts of their genome which may affect pathogenesis. Of particular interest to us are the mutations present in the accessory genes, which have been shown to contribute to pathogenesis in the host through innate immune signaling, among other effects on host machinery. To examine the effects of accessory protein mutations and other non-spike mutations on SARS-CoV-2 pathogenesis, we synthesized viruses where the WA1 Spike is replaced by each variant spike genes in a SARS-CoV-2/WA-1 infectious clone. We then characterized the *in vitro* and *in vivo* replication of these viruses and compared them to the full variant viruses. Our work has revealed that non-spike mutations in variants can contribute to replication of SARS-CoV-2 and pathogenesis in the host and can lead to attenuating phenotypes in circulating variants of concern. This work suggests that while Spike mutations may enhance receptor binding and entry into cells, mutations in accessory proteins may lead to less clinical disease, extended time toward knowing an infection exists in a person and thus increased time for transmission to occur.

**Significance:** A hallmark of the COVID19 pandemic has been the emergence of SARS-CoV-2 variants that have increased transmission and immune evasion. Each variant has a set of mutations that can be tracked by sequencing but little is known about their affect on pathogenesis. In this work we first identify accessory genes that are responsible for pathogenesis in vivo as well as identify the role of variant spike genes on replication and disease in mice. Isolating the role of Spike mutations in variants identifies the non-Spike mutations as key drivers of disease for each variant leading to the hypothesis that viral fitness depends on balancing increased Spike binding and immuno-evasion with attenuating phenotypes in other genes in the SARS-CoV-2 genome.

## Introduction

In December of 2019, a cluster of viral pneumonia cases were observed in Wuhan, Hubei Province, China.^1^ The etiologic agent of this infection was found to be a novel coronavirus that we now call severe acute respiratory syndrome coronavirus 2 (SARS-CoV-2).^2^ By early 2020, the virus was rapidly spreading, leading to infections on all seven continents and in every country around the world. There have since been over 420 million cases and almost six million deaths from this virus.^3^ Despite the rapid development and deployment of vaccines, the pandemic persists.

SARS-CoV-2 is a single-stranded positive-sense RNA virus that is 79% identical in sequence to SARS-CoV-1, the virus responsible for localized epidemic outbreaks beginning in February 2003.^4^ The genome of this and other beta coronaviruses is composed of Open Reading Frames (ORFs) which are functionally divided between replicase proteins, structural proteins, and accessory proteins, the latter of which are unique to each CoV species. ^5,6^ From the 5’ to 3’ end, the virus encodes the replicase (ORF1a/b) and the four ORFS for the structural proteins spike (S), envelope (E), membrane (M), and nucleocapsid (N). The replicase is responsible for encoding sixteen nonstructural proteins that compose the replicative machinery of the virus. Additionally, interspersed with the structural proteins at the 3’ end of the genome is a variety of accessory ORFs. The accessory ORFs encode proteins which are not essential for viral replication *in vitro* but generally contribute to viral pathogenesis. The accessory ORFs of SARS-CoV-2 are very similar to those of SARS-CoV-1, and many of the functions of these ORFs have been inferred based on the previously identified functions of the SARS-CoV-1 accessory ORFs. ^5^

The functions of the accessory ORFs of SARS-CoV-2 involve modulation of several different host pathways including antagonism of the innate immune response. For example, SARS-CoV-2 ORF3b has been shown to antagonize interferon signaling and ORF7a has been shown to interfere with the interferon-stimulated gene (ISG) BST2.^7,8,9^ SARS-CoV-2 ORF6 also participates in this antagonism of the innate immune response as it has been shown to antagonize the IFN-induced nuclear translocation of STAT1 which results in the reduced expression of ISGs.^10^

The continuation of the COVID-19 pandemic is largely due to the emergence of mutated strains or “variants” of SARS-CoV-2. The variants differ most notably in the sequence of their spike proteins, which bind to the receptor angiotensin-converting enzyme 2 (ACE2) to allow for internalization of the virus. As the spike protein is the immunodominant antigen, the emergence of variants has raised concerns regarding the breadth of protection of the SARS-CoV-2 vaccines. However, it is important to note that many of the variants of SARS-CoV-2 possess mutations in one or more of the accessory proteins. The impact of such mutations outside of the spike protein on pathogenesis of these variants remains understudied.

To elucidate the role of the accessory proteins of SARS-CoV-2 in pathogenesis, we developed a synthetic genomics assembly approach based on transformation-associated recombination (TAR) in yeast for the creation of infectious clones of SARS-CoV-2.^11–14^ We then synthesized deletion viruses of ORFs 3a/3b, 6, 7a/7b, and 8 in the prototype SARS-CoV-2 (isolate USA/-WA1/2020 or WA-1) strain of SARS-CoV-2. We then investigated the replicative fitness of these viruses *in vitro* before proceeding to characterization of the effect of these accessory deletions on pathogenesis in a murine model. To begin to characterize the impact of naturally occurring accessory mutations found in the variants on pathogenesis of SARS-CoV-2, we developed recombinant variant spike proteins in the WA-1 backbone (B.1.1.7, B.1.351, and P.1). We then compared the replicative fitness *in vitro* of these recombinant viruses to the parent variants and characterized differences in pathogenesis in a mouse model.

## Methods

### Infectious Clone Construction and Rescue

#### TAR-cloning SARS-CoV-2 DNA fragments

To clone each SARS-CoV-2 DNA fragment, each of the target PCR amplicons of the genome was cloned into a TAR vector backbone for replication in budding yeast and *Escherichia coli*. TAR vectors consist of the vector backbone flanked by restriction sites to release the genomic region and homologous “hooks” of sequence. To assemble DNA fragment clones, the TAR vectors were PCR amplified from pCC1BAC-his3 with KOD Xtreme Hot Start DNA polymerase (*Millipore*, Burlington, MA) using the construction primers (labeled “Con”, Table S1).^15^ 40 bp of homology and an I-SceI restriction site were introduced by the Con primers to each end of the vector backbone. Each DNA fragment was amplified with KOD Xtreme Hot Start DNA polymerase (*Millipore*,Burlington, MA) using cDNA from SARS-CoV-2/human/USA/WA-CDC-WA1/2020 and contains 80 bp of homology to its adjacent fragments for assembly. PCR products were digested with DpnI (*New England Biolabs*, Ipswich, MA) before transformation as per the manufacturer’s directions.

The SARS-CoV-2 WA1 genome was cloned as 7 individual DNA fragments: 1a-1, 1a-2, 1a-3, 1b-1, 1b-2, S and AP. 50 fmol of each WA1 PCR amplicon was transformed with 15 fmol of the appropriate amplified YCpBAC vector into *Saccharomyces cerevisiae* strain VL6-48N spheroplasts (*MATα, his3-Δ200, trp1-Δ1, ura3-Δ1, lys2, ade2–101, met14, cir°*) as described in.^13–15^ Transformants were patched on synthetic dropout medium plates without histidine (SD-his). After sufficient growth, yeast cells were lysed by incubation in 25 mM NaOH at 95°C for 30 min. Detection PCR amplification, with Q5 polymerase (*New England Biolabs*, Ipswich, MA), lysed cell supernatant as template, and detection primers (RCO493 + “Det-F” or RCO495 + “Det-R” primers, Table S2), was used to confirm correct junctions between the fragment and vector. Sanger sequencing of PCR products (*GeneWiz, Inc*., South Plainfield, NJ) across the SARS-CoV-2 regions was used to verify the DNA fragment clone. DNA of sequenced-validated clones was extracted from yeast patches and electroporated into *E. coli* DH10B (Thermo Fisher).^13^ *E. coli* transformants were screened by colony PCR amplification for correct junction sizes with the appropriate “F” and “R” detection primers (Table S2). Plasmid DNAs from the DNA fragment clones were isolated from *E. coli* by the Purelink HiPure Plasmid Midiprep Kit (*Thermo Fisher*, Waltham, MA) and modified or used for full-length genome assembly.

#### DNA fragment modifications

*In vitro* CRISPR-Cas9 digestion and TAR assembly were used to make accessory ORF deletions in the AP DNA fragment. Cas9 target sites near the desired mutant coordinate were identified using Benchling’s CRISPR guide RNA design tool (www.benchling.com/crispr/). Single guide RNA (sgRNA) was transcribed from a PCR amplicon, generated as described in PMID: 28928148, containing 18 bp of specific target sequence upstream of the PAM sequence. The AP DNA fragment was digested *in vitro* using Cas9 nuclease, *Streptococcus pyogenes* (*New England Biolabs*, Ipswich, MA), as described previously ^13^ An oligonucleotide containing 40 bases of homology to either side of the target accessory ORF was TAR-assembled with each appropriate Cas9-digested AP DNA fragment in yeast by spheroplast transformation. Yeast transformants were screened for the desired mutation by Sanger sequencing of PCR amplicons (*GeneWiz, Inc*., South Plainfield, NJ) using lysed yeast culture supernatant as template. DNA was then extracted from yeast, transformed into *E. coli*,screened for correctness, and isolated from *E. coli* as with DNA fragment clones described above.

SARS-CoV-2 variant spike genes were generated as described previously.^25^ Briefly, to generate SARS-CoV-2 variant spike genes, short fragments were amplified with Platinum SuperFi II DNA polymerase (*Thermo Fisher*, Waltham, MA) using the wild-type S DNA fragment clone as template. Variant mutations (**Table S6**) were introduced by primers into each amplicon, which had 30-35 bp homology at each end between the adjacent fragments. Amplicons were digested with DpnI (*New England Biolabs*, Ipswich, MA) to remove template DNA and purified with a Qiagen PCR purification kit. 50 fmol of each amplicon and 15 fmol of YCpBAC vector were assembled using a standard Gibson assembly reaction (*New England Biolabs*, Ipswich, MA), transformed into *E. coli* DH10B competent cells (*Thermo Fisher*, Waltham, MA), and plated on LB medium with 12.5 mg/ml chloramphenicol. Assembled plasmids were confirmed to have the desired mutations by colony PCR amplification and Sanger sequencing (*GeneWiz, Inc*., South Plainfield, NJ), and then isolated from *E. coli* using the Purelink HiPure Plasmid Midiprep Kit (*Thermo Fisher*, Waltham, MA). **Table S3** lists the primers used for the construction of the variant S DNA fragments.

#### Full-length genome assembly

The TAR vector for assembly of the full-length genome was amplified from pCC1BAC-ura3 using primers ConCMVpR and ConBGHtermF with KOD Xtreme Hot Start DNA polymerase (*Millipore*, Burlington, MA).^15^ The CMV promoter was amplified from HCMV Toledo genomic DNA using primers CMVpromF and CMVpromR. Hepatitis delta virus ribozyme (HDV Rz) with bovine growth hormone (BGH) terminator fragment was obtained as a gBlock (IDT), Hu1-34. A fragment to introduce a BamHI restriction site and a fragment with a polyA region were synthesized by IDT (Pcmv-BamHI fix and PA35-HDV fix, Table S4). Each fragment has 40 bp of homology at each end to its adjacent fragments. To generate the TAR vector for the complete genome, the pCC1-ura3 amplicon, the CMV amplicon, the BamHI fragment, the polyA fragment, and the HDVR+BGH region were assembled into circular DNA in yeast by TAR.

SARS-CoV-2 DNA fragments, isolated from *E. coli*, were digested with I-SceI (*New England Biolabs*, Ipswich, Ma) to release the overlapping DNA fragments from the vector whereas the full-length TAR vector was linearized by BamHI (*New England Biolabs*, Ipswich, MA) digestion. 50 fmol of each fragment and 15 fmol of full-length genome TAR vector were mixed with yeast spheroplasts for TAR-assembly.^12,13^ Transformants were patched on SD-URA plates, and positive clones were screened by PCR amplification with the detection primers listed in **Table S5**. The subsequent DNA isolation from yeast, transformation into and isolation from *E. coli* were performed as described for the SARS-CoV-2 DNA fragment clones.

#### Virus Reconstitution

24 hours prior to transfection, 5e4 VeroE6 cells (*ATCC*,Manassas, VA) were plated per well in 1mL of VeroE6 media (DMEM (*Quality Biological*, Gaithersburg, MD), 10% FBS (*Gibco*, Waltham, MA), 1% Penicillin-Streptomycin (*Gemini Bio Products*, Sacramento, CA), 1% L-Glutamine (*Gibco*, Waltham, MA)). For transfection, 5μg of the infectious clone and 100ng of a SARS-CoV-2 WA-1 nucleoprotein expression plasmid were diluted in 100μL of OptiMEM (*Gibco*, Waltham, MA). 3μL of TransIT-2020 (*Mirus Bio*, Madison, WI) was added and the reactions were incubated for 30 minutes prior to addition to the cells in the BSL-3. Cells were checked for cytopathic effect (CPE) 72-96 hours after transfection and the supernatant collected for plaque purification. For plaque purification, 6e5 VeroE6 cells were plated in a 6-well plate in 2mL of VeroE6 media 24 hours prior to infection. 25μL of this supernatant was then serial diluted 1:10 in DMEM and 200μL of this supernatant was added to the VeroE6 cells. The cells were rocked every 15 minutes for 1 hour at 37°C prior to overlay with 2mL of a solid agarose overlay (EMEM (*Quality Biological*, Gaithersburg, MD), 10% FBS, 1% Penicillin-Streptomycin, 1% L-Glutamine, 0.4% w/v SeaKem agarose (*Lonza Biosciences*,Morrisville, NC). Cells were incubated for 72 hours at 37°C and 5% CO_2_ and individual plaques were picked and then transferred to a well of a 6-well plate with 4e5 VeroE6 cells in 3mL VeroE6 media. After 48 hours, successful plaque picks were assessed by presence of CPE. 1mL of a well showing CPE was transferred to a T175 with 8e6 VeroE6 cells in 30mL of VeroE6 media and the virus stock was collected 48 and 72 hours after. The stocks were then titered by plaque assay.

### Titering of Virus Stocks, Growth Curve Samples, Tissue Homogenates by Plaque Assay

The day prior to infection, 2e5 VeroE6 cells were seeded per well in a 12-well plate in 1mL of VeroE6 media. Tissue samples were thawed and homogenized with 1mm beads in an Omni Bead ruptor (*Omni International Inc*., Kennesaw, GA) and then spun down at 21,000xg for 2 minutes. A 6-point dilution curve was prepared by serial diluting 25μL of sample 1:10 in 225μL DMEM. 200μL of each dilution was then added to the cells and the plates were rocked every 15 minutes for 1 hour at 37°C. After 1hr, 2mL of a semi-solid agarose overlay was added to each well (DMEM, 4%FBS, 0.06% UltraPure agarose (*Invitrogen*, Carlsbad, CA). After 72 hours at 37°C and 5% CO_2_, plates were fixed in 2% PFA for 20 minutes, stained with 0.5mL of 0.05% Crystal Violet and 20% EtOH, and washed 2x with H_2_O prior to counting of plaques. The titer was then calculated. For tissue homogenates, this titer was multiplied by 40 based on the average tissue sample weight being 25mg.

### Growth Curve Infection and Sample Processing

The day prior to infection, 1.5e5 VeroE6 cells were seeded in a 12-well plate in 1.5mL of VeroE6 media. The day of infection, cells were washed with 500μL of DMEM and the volume of virus needed for an M.O.I. of 0.01 was diluted in 100μL DMEM and added to a well in triplicate. The plates were rocked every 15 minutes for 1hr at 37°C. After 1hr, the inoculum was removed, the cells were washed with 500μL of VeroE6 media and then 1.5mL of VeroE6 media was added to each well. 300μL of this supernatant was taken as the 0hr timepoint and replaced with fresh media. 300μL of supernatant was pulled and replaced with fresh media at 0hr, 6hr, 24hr, 48hr, 72hr, and 96hr timepoints. This supernatant was then titered.

### Infection of BALB/c and hACE2/k18 Mice

All animals were cared for according to the standards set forth by the Institutional Animal Care and Use Committee at the University of Maryland-Baltimore. On Day 0, 12-week old BALB/c (*Charles River Laboratories*, Wilmington, MA) and 12-week old hACE2 transgenic K18 mice (K18-hACE2) (*Jackson Labs*, Bar Harbor, ME) were anesthetized interperitoneally with 50μL ketamine(1.3mg/mouse)/ xylazine (0.38mg/mouse). The BALB/c mice were then inoculated with 1e4 PFU of each virus in 50μL PBS. The K18-hACE2 mice were inoculated with 1e3 PFU of each virus in 50μL PBS. The mock infected mice received 50μL PBS only. Mice were then weighed every day until the end of the experiment. Mice were euthanized with isoflurane on Day 2 and Day 4. From each mouse, the left lung was collected in PFA for histology and the right lung was split in half with one half placed in PBS for titer and one half placed in TRIzol for RNA extraction. For the K18-hACE2 mice, the brain was also collected, with half of the brain in PBS for titering and the other half in TRIzol for RNA extraction.

### RT-qPCR of Tissue Homogenates

Samples were homogenized in an Omni Bead ruptor in TRIzol and then the RNA was extracted from 300μL of each sample using the Direct-zol RNA miniprep kit (*Zymo Research*, Irvine, CA). 2μL of isolated RNA from each sample was then converted to cDNA using the RevertAID first strand cDNA synthesis kit (*Thermo Fisher*, Waltham, MA) in a 20μL volume. For qPCR for SARS2 Rdrp, 20μL reactions were prepared using 2μL cDNA, 1μL of 10mM Rdrp Forward primer (10006860, *Integrated DNA Technologies*, Coralville, IA), 1μL of Rdrp Reverse primer (10006881, *Integrated DNA Technologies*, Coralville, IA), and 10μL of 2x SYBR Green (*Thermo Fisher*, Waltham, MA). The reactions were then run on a 7500 Fast Dx Real-Time PCR Instrument (4357362R, *Applied Biosystems*, Waltham, MA). For qPCR for murine GAPDH, 20μL reactions were prepared using 2μL cDNA, 1μL of a 20x murine GAPDH primer (MM.pt.39a.1, *Integrated DNA Technologies*, Coralville, IA), and 10μL of 2x SYBR Green. The reactions were then run on a QuantStudio 5 Real-Time PCR Instrument (A28133, *Applied Biosystems*, Waltham, MA).

### H/E Staining of Lungs and Pathological Scoring

Lungs were scored in a blinded fashion with a 0-5 score given, 0 being no inflammation and 5 being the highest degree of inflammation. Interstitial inflammation and peribronchiolar inflammation were scored separately then scores were averaged for the overall inflammation score.

### Cytokine Arrays

The concentration of lung RNA was quantified using a Nanodrop (NanoVue Plus). 400ng of RNA was converted to cDNA using the Qiagen RT^2^ First Strand Kit (330404, *Qiagen*, Hilden, Germany). The cDNA was analyzed with the Qiagen RT^2^ Mouse Cytokines and Chemokines array (PAMM-150Z, *Qiagen*, Hilden, Germany). Reactions were run on a QuantStudio 5 Real-Time PCR Instrument (A28133, *Applied Biosystems*, Waltham, MA). The results were analyzed with the Qiagen analysis spreadsheet provided with the kit.

### Statistical Analysis

All statistical analyses were carried out using the GraphPad Prism software. The statistical tests run were unpaired t-tests assuming unequal variances. The cutoff value used to determine significance was p≤0.05.

### Biosafety approval

All virus experiments and recombinant virus creation was approved by the Institutional Biosafety Committee at The University of Maryland, Baltimore. At their request, the P.1 S in WA-1 virus has been destroyed due to it increased in vivo pathogenesis compared to the clinical P.1 strain.

## Results

### Infectious Clone System

To enable rapid and combinatorial modification of SARS-CoV-2, we established a yeast-based assembly approach using transformation-associated recombination (TAR) to assemble a complete genome from overlapping DNA fragments. This approach is based on one that we previously used to engineer human herpesviruses.^13,14^ It involves first, generating a set of sequence-validated DNA fragments that encompass a complete genome which can then be modified individually and in parallel to obtain DNA fragments containing the desired changes. The modified fragments can be mix and matched with unmodified DNA fragments for assembly into complete genomes to generate the desired mutant viruses. For SARS-CoV-2, the genome was deconstructed into 7 individual DNA fragments: 3 for ORF1a (1a-1 to 1a-3), 2 for ORF1b (1b-1 and 1b-2), 1 for Spike (S) and 1 for the accessory protein 3’ genome region (AP) [**Figure 1A**]. Each individual fragment was TAR-cloned by transforming into yeast each target fragment amplified by PCR using SARS-CoV-2 WA1 cDNA as template with primer pairs listed in **Table S1** together with a PCR-amplified YCpBAC vector that contains each set of appropriate homologous sequences at either end (**Table S1**). Each individual DNA fragment was confirmed by junction PCR amplification using a primer homologous to the YCpBAC vector and a primer homologous to the target DNA fragment (**Figure S1, Table S2**). Positive yeast clones were then transformed into *Escherichia coli* to enable isolation of large amounts of DNA and again confirmed by junction PCR amplification (**Figure S1**). Finally, each individual DNA fragment, which was validated by Sanger sequencing, is flanked by I-SceI sites and contains 80 base pairs of homologous sequence to its adjacent fragments. Deletion of the accessory genes were performed at the DNA fragment level (AP) by cleaving with Cas9 protein *in vitro* guided by guide RNAs specific to each accessory gene and then transforming into yeast each cleaved DNA fragment together with the appropriate “fix” that contains 40 bases of homology to either side of the coding region of the target accessory gene (**Table S3**). Positive clones were identified by PCR amplification across the target gene and then transformed into *E. coli* for large quantity isolation of DNA (**Figure S2**). The Spike ORFs from select SARS-CoV-2 variants were generated by rebuilding the WA1 S DNA fragment by Gibson assembly in *E. coli* from PCR amplified products using primers to introduce the changes where needed and the WA1 S DNA fragment as template (**Table S3**). Each of the SARS-CoV-2 S variant generated was validated by Sanger sequencing. Finally, to generate the different full-length genomes by TAR assembly, the appropriate sets of DNA fragments (1a-1 to AP) were cleaved with I-SceI to separate the SARS-CoV-2 fragment from the YCpBAC vector and then transformed into yeast together with a YCpBAC vector that contains a CMV promoter and homology to DNA fragment 1a-1 at one end and homology to DNA fragment AP, Hepatitis delta virus ribozyme sequence as well as bovine growth hormone terminator at the other end (**Figure 1A, Table S4**). Correct assembly of the fulllength genomes were confirmed by junction PCR amplification (**Figure 1B**) using primer pairs that span the SARS-CoV-2 genome (**Table S5**). Positive clones were transferred to *E. coli* for isolation of SARS-CoV-2 full-length genome DNA.

**Figure 1.**
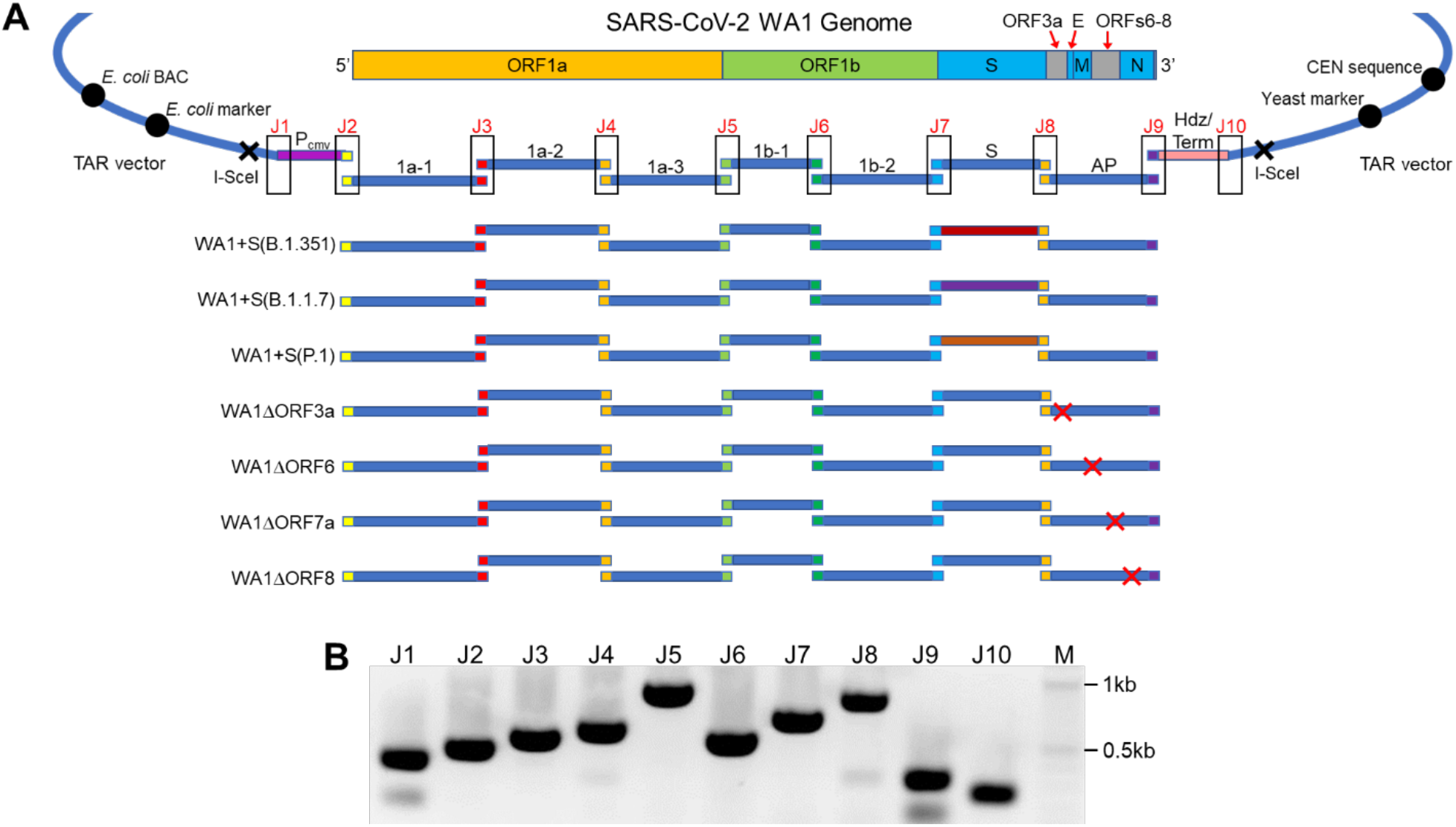
Assembly of infectious clone genomes of SARS-CoV-2. A. The genome of WA1 was assembled from sequence-validated overlapping (colored ends) DNA fragments (1a-1, 1a-2, 1a-3, 1b-1, 1b-2, S, AP; blue lines) by TAR in yeast. The infectious clone genomes can be maintained in yeast and *E. coli*-by a YCpBAC vector. The infectious clone genome, which is flanked by I-SceI sites, is driven by a CMV promoter (P_CMV_) and has a hepatitis delta virus ribozyme sequence (Hdz) as well as a bovine growth hormone terminator (Term) at the 3’ end of the genome. SARS-CoV-2 WA1 genomes containing either Spike variants or accessory ORF deletions were assembled from a mix of unmodified and appropriate modified DNA fragments. B. PCR amplifications of assembly junctions (J1-10) to confirm a full-length genome in *E. coli*. M, 2-log marker.

Following confirmation of the successful synthesis of each full-length genome clone, VeroE6 cells were transfected with each isolated YCpBAC. 72-96 hours later, the cells were monitored for CPE, supernatant was collected. The virus was then plaque-purified and stocked. The accessory ORF deletion viruses of WA-1 were successfully recovered and sequence-verified prior to use in experiments.

### Replication of SARS-CoV-2 deletion viruses in cells

The growth of the wildtype SARS-CoV-2 (WA-1) and the accessory ORF deletion series was analyzed. To evaluate the replicative kinetics of the deletion viruses *in vitro*, VeroE6 cells were infected with an M.O.I. of 0.01 of WA-1, WA-1ΔORF3a/b, WA-1ΔORF6, WA-1ΔORF7a/b, and WA-1ΔORF8. Supernatant was titered at 0, 6, 24, 48, 72, and 96 hours post infection by plaque assay. Compared to the wildtype WA-1 virus, WA-1ΔORF3a/b was the only accessory gene deletion virus to show an attenuated phenotype, demonstrating significantly attenuated growth at 48 and 72 hours (**Figure 2A** p=0.0027 and p=0.0014).

**Figure 2.**
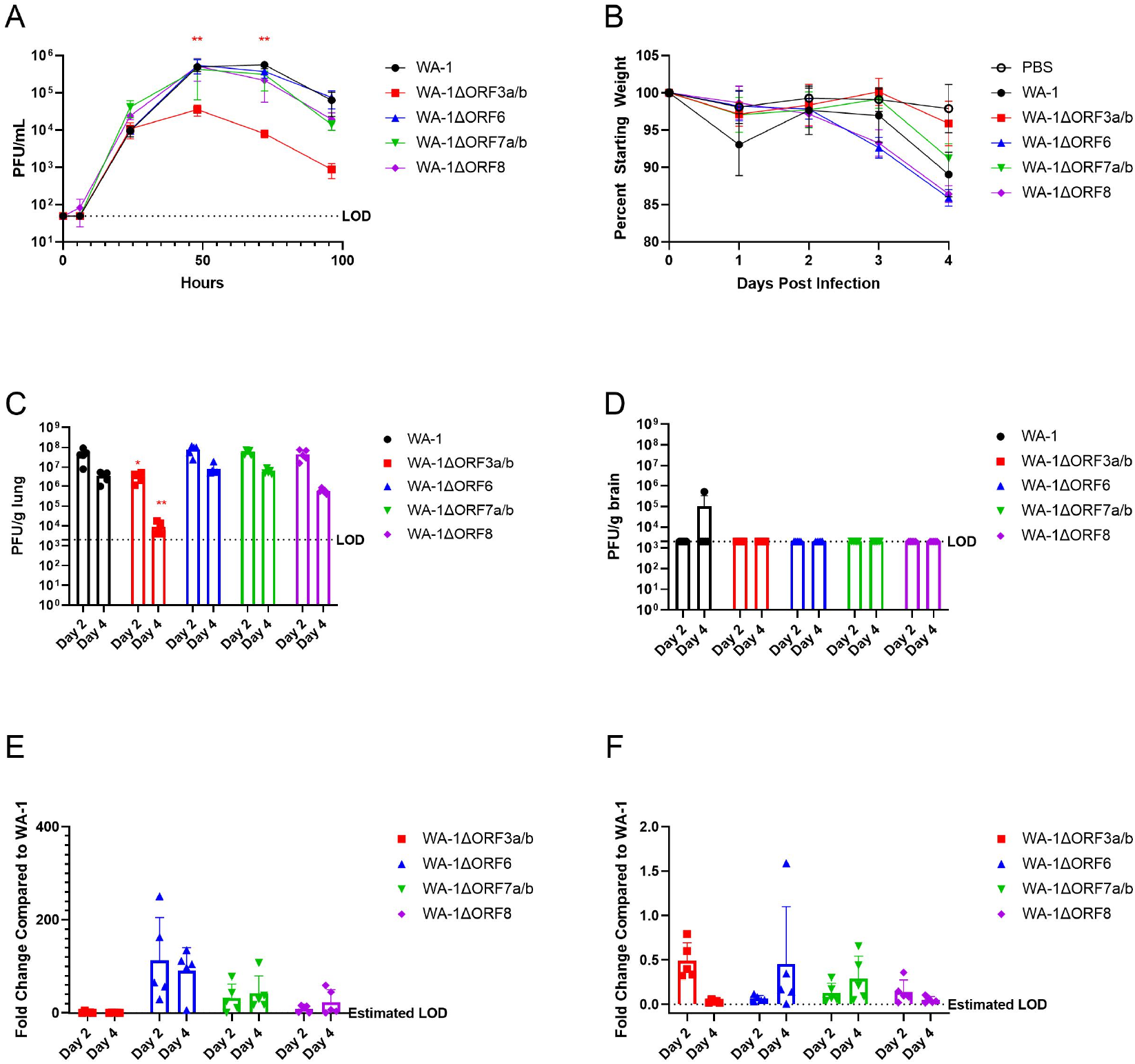
WA-1 accessory deletion viruses in 12-week old K18-hACE2 mice. A. Supernatant titers of VeroE6 cells infected with an M.O.I. of 0.01 of accessory deletion viruses with supernatant pulled at 0, 6, 24, 48, 72, and 96 hours and titered by plaque assay. B. Percent starting weight on days 0-4 of K18-hACE2 mice infected with 1e3 pfu of each accessory deletion virus. C. Lung viral titers of mice euthanized on D2 and D4 by pfu/g lung. D. Brain viral titers of mice euthanized on D2 and D4 by pfu/g brain. E. Lung viral loads of mice euthanized on D2 and D4 by qPCR for Rdrp. F. Brain viral loads of mice euthanized on D2 and D4 by qPCR for Rdrp. (*= pfu ≤ 0.05, **=pfu ≤ 0.005, ***= pfu ≤ 0.0005)

### WA-1 Accessory Deletion Viruses in K18-hACE2 mice

We sought to identify the role of the accessory proteins in SARS-CoV-2 pathogenesis using a mouse model of COVID-19. C57BL/6 K18/hACE2 (K18-hACE2) are highly permissive to SARS-CoV-2 due to the widespread expression of human ACE2 (hACE2). This hACE2 is expressed under the control of the keratin 18 (K18) promoter, leading to hACE2 expression in almost all cells of the mouse. Previous reports have shown this mouse strain to be permissive to wildtype SARS-CoV-2 infection without the need for any adaptations or mutations in the Spike protein. We utilized these mice in the comparison of wildtype and deletion strains so that no alteractions to the Spike protein would be needed for replication. To evaluate the role of each accessory protein, 12-week old K18-hACE2 mice were infected with 1e3 pfu of each virus, weighed daily, and euthanized at 2 and 4 days post-infection (dpi) for collection of tissue samples. Wildtype WA-1 displayed ~12% weight loss through 4 days of infection, with similar weight loss for WA-1ΔORF6, WA-1ΔORF7a/b and WA-1ΔORF8 viruses (**Figure 2B**). However, infection with WA-1ΔORF3a/b resulted in an attenuated weight loss phenotype in the mice, with these mice losing less than 4% of their starting weight (**Figure 2B**).

Lung titers for wildtype and deletion viruses were analyzed at day 2 and 4 post infection to determine if there were differences in replicative fitness between the mutants and WA-1. Lung titers for WA-1 reached 7e7 and 8e6 pfu/g lung at day 2 and 4, respectively (**Figure 2C**). The mutant viruses WA-1ΔORF6, WA-1ΔORF7a/b and WA-1ΔORF8 had similar lung titers to WA-1 at day 2. This was also found with infection of WA-1ΔORF6 and WA-1ΔORF7a/b at day 4. WA-1ΔORF8 did exhibit a one log reduction in lung titer at day 4, although this reduction was not significant (**Figure 2C**). Consistent with the observed attenuation in weight loss, mice infected with WA-1ΔORF3a/b had significantly attenuated lung titers at both day 2 and 4 with one log lower lung titer at day 2 (p=0.012) and three logs lower titer at day 4 post infection (p=0.0030) compared to WA-1 (**Figure 2C**).

Lung viral RNA was analyzed for WA-1 and each mutant virus at day 2 and 4 post infection. The levels for viral RNA for each of the deletion viruses were compared to the RNA levels by fold change in the lungs of the mice infected with the wildtype virus. Although none of the fold changes were significant, mice infected with the deletion viruses for ORF6, ORF7a/b, and ORF8 showed higher viral RNA levels compared to WA-1 while mice infected with the ORF3a/b deletion virus showed slightly lower lung RNA levels compared to WA-1 (**Figure 2E**).

Due to the over-expression of hACE2 in neural tissues in the K18-hACE2 mice, SARS-CoV-2 can invade and replicate in the brain of these mice. We therefore assessed the replication of WA-1 and the accessory deletion viruses in brain tissue as well. For all of the accessory deletion viruses, there were undetectable live virus titers in the brain at day 2 and day 4 postinfection (**Figure 2D**). Despite there being little virus in the brain by plaque assay, there was viral RNA detected in the brain in levels that were not significantly different between the accessory deletions when compared to WA-1 (**Figure 2F**).

Lungs were fixed and stained with H&E to determine whether inflammation and lung damage were different between the wildtype WA-1 and deletion mutants. They were scored in a blinded fashion with a 0-5 score given, 0 being no inflammation and 5 being the highest degree of inflammation. Interstitial inflammation and peribronchiolar inflammation were scored separately then scores were averaged for the overall inflammation score. Despite the attenuation of weight loss with infection with WA-1ΔORF3a/b, there is no significant difference in lung pathology in the mice when compared to WA-1 (**Figure 3A, 3B**). Across the accessory deletion panel, there were no significant differences in lung pathology seen with any of the accessory deletion viruses except for an unexpected increase in lung pathology scores for mice infected with WA-1ΔORF8 at day 4 (p=0.0037, **Figure 3A, 3B**). The increased pathology of WA-1ΔORF8 is intriguing because of the reported immunomodulatory role of the secreted ORF8 protein in pathogenesis. This finding is being studied in future experiments.

**Figure 3.**
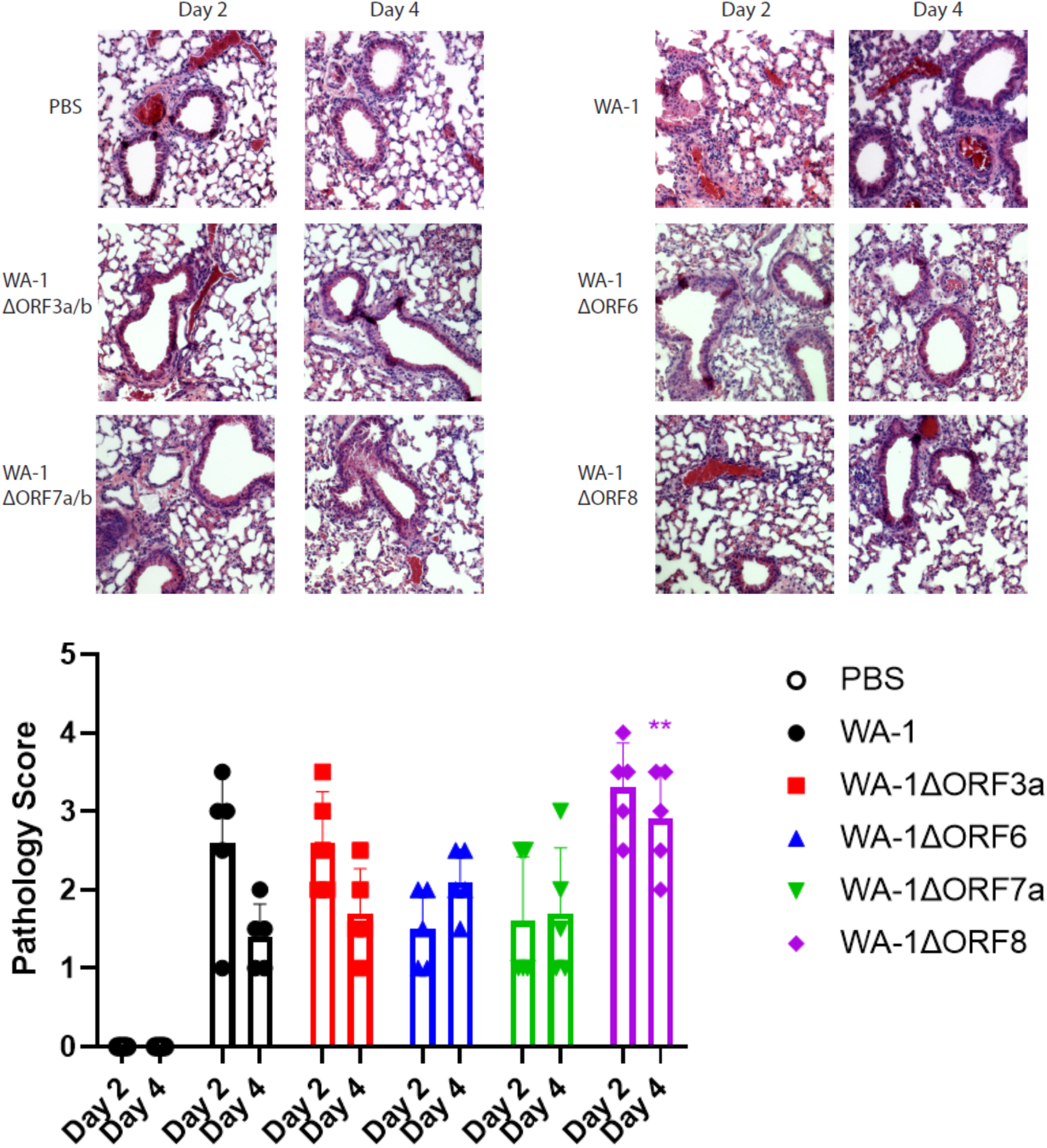
Lung pathology of 12-week old K18-hACE2 mice infected with accessory deletion viruses of WA-1. A. H/E stained sections of the lungs. B. Pathological scoring of the lungs.

### Creation of infectious clones of variant spikes in a WA-1 (icWA-1) background and analysis in cells

The Spike ORFs from selected SARS-CoV-2 variants were cloned into the icWA-1 clone to replace the WA-1 Spike ORF. The goal of these mutant viruses is to dissociate the effects of the variant Spike ORF from the other mutations in the genomes and to determine if differences exist in pathogenesis in the mouse model. This allows for the identification of whether the variant viruses replicate and have similar pathogenesis to a virus containing only the Spike mutations alone on the same background SARS-CoV-2 virus.

The Spike variant viruses were cloned, rescued, sequenced and analyzed for their replication in vitro and in vivo. Upon successful recovery and sequence verification of the variant spikes in WA-1 viruses, the replication of these viruses was compared to the parent variants at an M.O.I. of 0.01 in VeroE6 cells. The variant spike and parental variant pairs, B.1.351 and B.1.351 S in WA-1 and B.1.1.7 and B.1.1.7 S in WA-1, showed no difference in supernatant viral titers at all timepoints tested (**Figure 4A**). The supernatant titer for P.1 S in WA-1 was significantly lower than the parent variant P.1 at only the 72hr timepoint (**Figure 4A**, p=0.0015).

**Figure 4.**
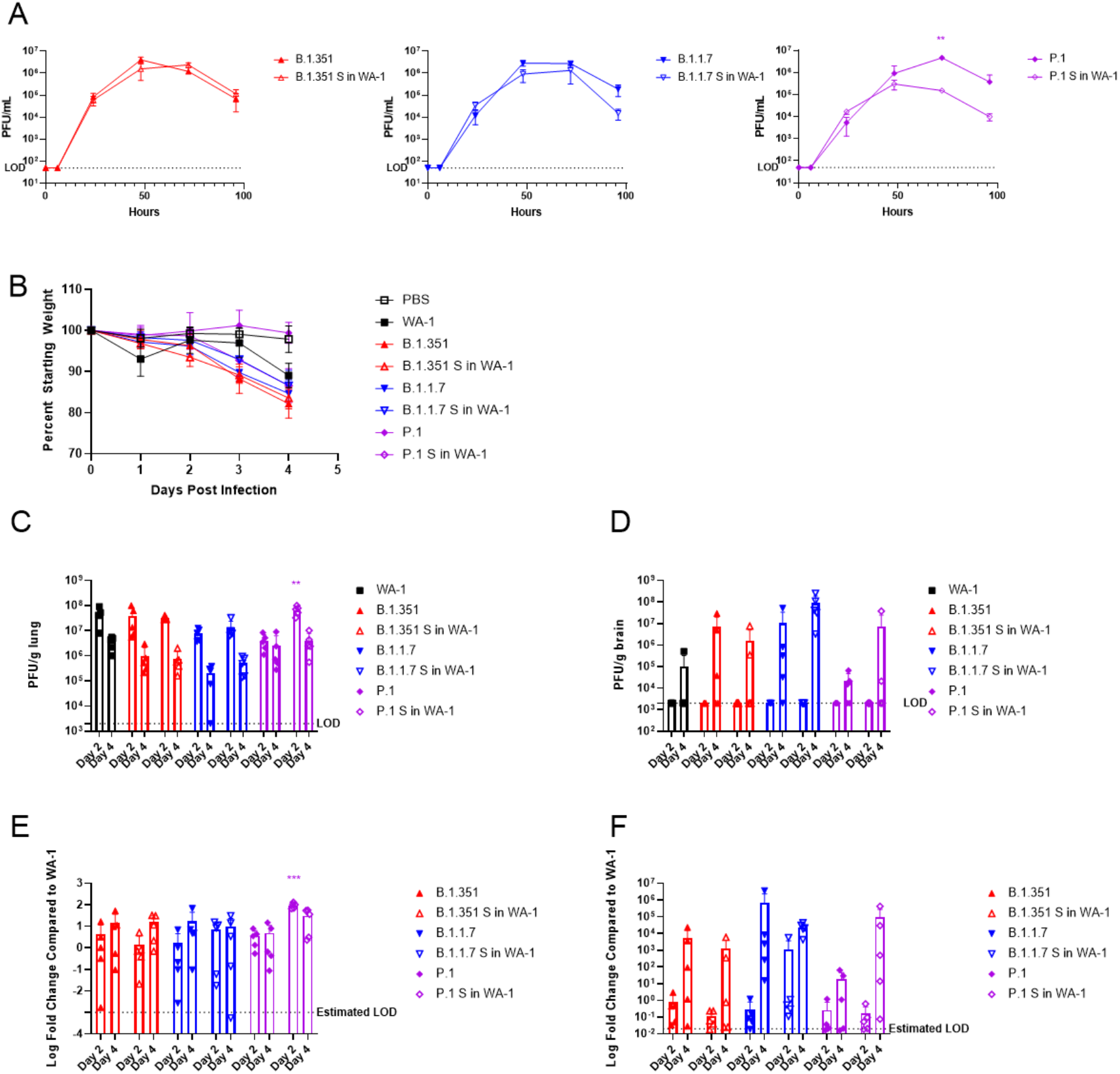
Variant spikes in WA-1 viruses in 12-week old K18-hACE2 mice. A. Supernatant titers of VeroE6 cells infected with an M.O.I. of 0.01 of variant spikes in WA-1 and parent variant viruses with supernatant pulled at 0, 6, 24, 48. 72, and 96 hours and titered by plaque assay. B. Percent starting weight on days 0-4 of K18-hACE2 mice infected with 1e3 pfu of each variant spike in WA-1 virus or the parent variant virus. C. Lung viral titers of mice euthanized on D2 and D4 by pfu/g lung. D. Brain viral titers of mice euthanized on D2 and D4 by pfu/g brain. E. Lung viral loads of mice euthanized on D2 and D4 by qPCR for Rdrp. F. Brain viral loads of mice euthanized on D2 and D4 by qPCR for Rdrp.

### Variant spike in WA-1 viruses compared to variant clinical isolates in K18-hACE2 mice

The variant Spike viruses were then tested to determine if their pathogenesis in mice was different than the parent clinical isolates that have the Spike mutations and the full composition of non-spike mutations. To investigate differences in pathogenesis *in vivo*, 12-week old K18-hACE2 mice were infected with 1e3 pfu of each virus. Mice were weighed every day and were euthanized at either day 2 or day 4 post-infection for collection of lung tissue and brain tissue. Reflective of the *in vitro* results, mice infected with B.1.351 and mice infected with B.1.351 S in WA-1 displayed similar weight loss of around 12%. Mice infected with B.1.1.7 and mice infected with B.1.1.7 S in WA-1 also exhibited similar weight loss of around 10% (**Figure 4B**). However, mice infected with P.1 S in WA-1 showed a weight loss ~12% despite there being very minimal weight loss in mice infected with the P.1 parent variant (**Figure 4B**). The lung titers of the mice reflected the weight loss, with mice infected with B.1.351 S in WA-1 or B.1.17 S in WA-1 displaying similar lung titers of around 10^8^ on day 2 and 10^6^ on day 4 which was seen in the mice infected with the parent variants (**Figure 4C**). Mice infected with P.1 S in WA-1 had significantly higher lung titers at day 2 compared to mice infected with P.1, which lung titers being roughly one log higher at this time (**Figure 4C**, p=0.00057).

As with the accessory deletion viruses, viral titers in the brain were also analyzed. Although none of the infected mice had measurable titer in the brain at day 2, the brain titers for day 4 trended with the lung titer results, with B.1.351 and B.1.351 S in WA-1 and B.1.1.7 and B.1.1.7 S in WA-1 showing similar titers of ~10^7^ - 10^8^ at day 4 (**Figure 4D**). P.1 S in WA-1 showed 100X higher titer in the brain at day 4 compared to P.1, although this was not statistically significant (**Figure 4D**).

Viral RNA levels were quantified in lung and brain to compare to live virus titer results. The viral RNA levels in the lung and in the brain are reflective of the titers (**Figure 4E** and **Figure 4F**), with the only significant difference being between the day 2 lung RNA levels of P.1 and P.1 S in WA-1 (**Figure 4E**, p=0.00070).

Lung inflammation was also analyzed across clinical isolates and Spike mutant viruses. The lungs for the variant spike in WA-1 mice looked similar in terms of inflammation to the lungs of the mice infected with the parental variant viruses (**Figure 5A**). Although there were no significant differences in lung inflammation between the parent variant viruses and the variant spike in WA-1 viruses, the P.1 S in WA-1 lungs had higher lung pathology scores on both day 2 and day 4 when compared to the P.1 mice, which trends with both the titer and the weight loss data (**Figure 5B**).

**Figure 5.**
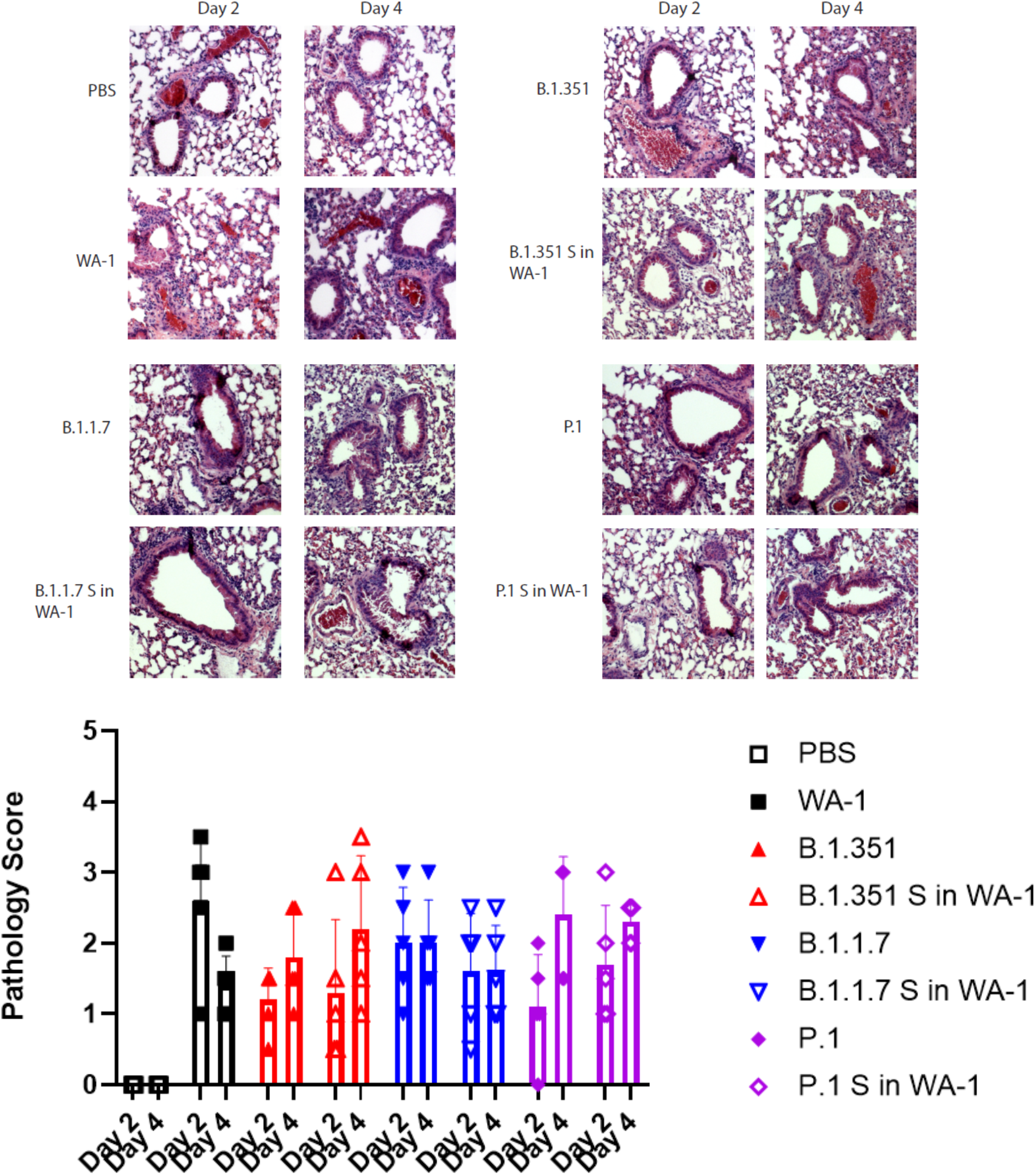
Lung pathology of 12-week old K18-hACE2 mice infected with variant spikes in WA-1 and parent variants. A. H/E stained lung sections. B. Pathological scoring of the lungs.

### Variant Spikes in WA-1 in BALB/c Mice

The wildtype SARS-CoV-2/WA-1 virus does not replicate in wildtype mice due to poor interactions with mouse ACE2. The N501Y mutation in Spike does however allow for replication in wildtype mice. Interestingly, the variants examined in this study do have the N501Y mutation allowing for comparison of pathogenesis between recombinant viruses variants and their parent strains in BALB/c mice. For this comparison, 12-week old BALB/c mice were infected with 1e4 pfu of each virus and euthanized at either Day 2 or Day 4 post-infection for analysis. As we observed with the K18-hACE2 mouse experiment, infection with P.1 S in WA-1 resulted in increased weight loss of around 5% compared to infection with P.1 which produced no weight loss. Both B.1.1.7 and B.1.1.7 S in WA-1 infection produced minimal weight loss in these mice. Unlike the results from the infection in K18-hACE2 mice, mice with B.1.351 S in WA-1 showed attenuated weight loss of around 7% compared to B.1.351, which produced roughly 12% weight loss (**Figure 6A**).

**Figure 6.**
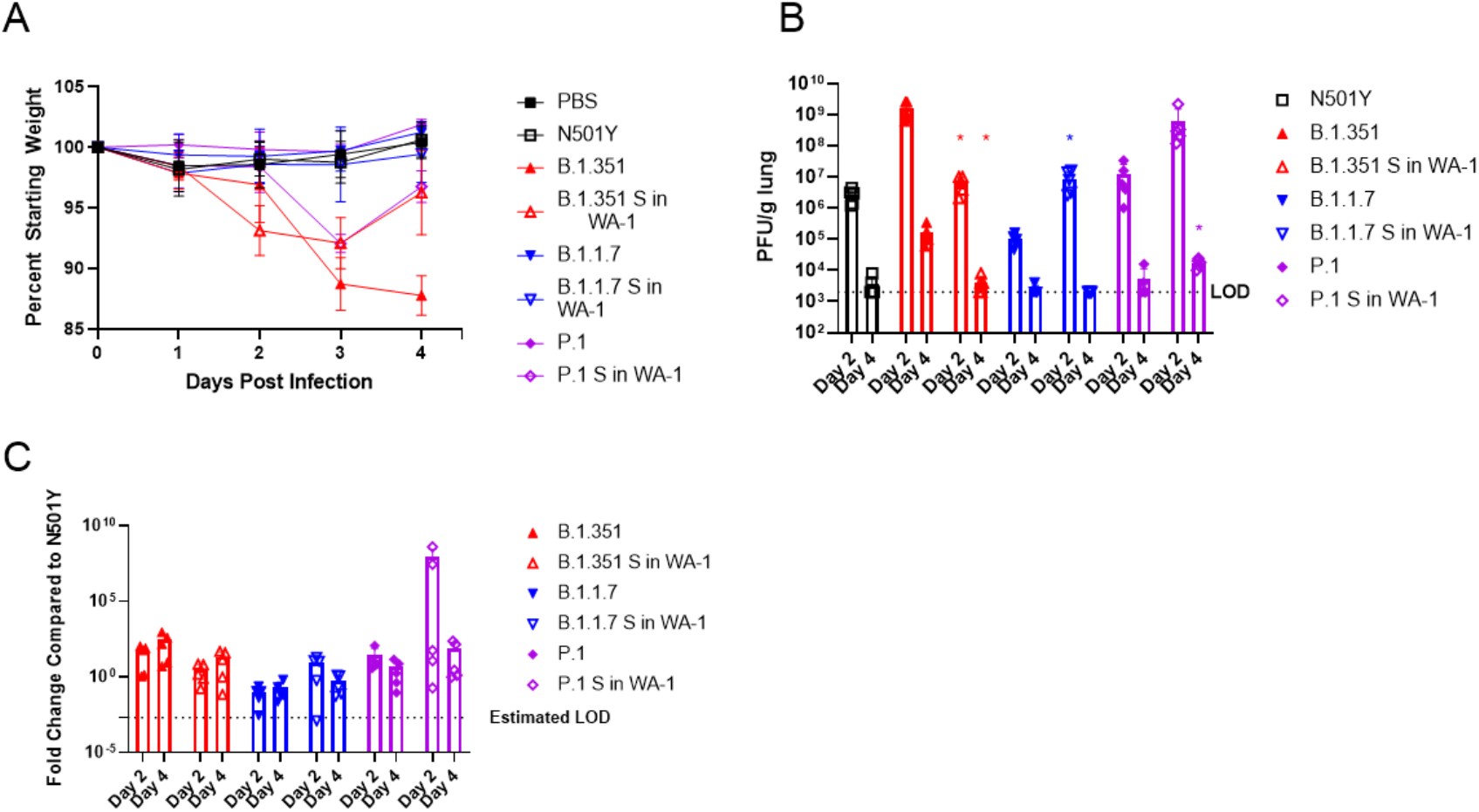
Variant spikes in WA-1 viruses in 12-week old BALB/c mice. A. Percent starting weight on day 0-4 of BALB/c mice infected with 1e4 pfu of each variant spike in WA-1 virus. B. Lung viral titers of mice euthanized on D2 and D4 by pfu/g lung. C. Lung viral titers of mice euthanized on D2 and D4 by qPCR for Rdrp.

The viral titers in the lung homogenates were mostly reflective of the weight loss findings, with infection with B.1.351 S in WA-1 producing significantly lower lung titers by about two logs at day 2 and day 4 as compared to infection with B.1.351 (**Figure 6B**, p=0.008 and p=0.014). Despite there being no differences in weight loss, B.1.1.7 S in WA-1 infection resulted in significantly higher lung titers by two logs at day 2 post infection (p=0.013) compared to B.1.1.7, with both viruses being nearly cleared by day 4 post infection (**Figure 6B**). Infection with P.1 S in WA-1 resulted in higher lung titers by two logs at day 2 and one log at day 4 as compared to infection with P.1, although only the day 4 titers differed significantly (**Figure 6B**, p=0.011).

The viral RNA levels in the lungs at these timepoints correlated to the viral titer levels, although none of these findings were statistically significant (**Figure 6C**). Lung pathology was also analyzed. The lungs of mice infected with B.1.1.7 S in WA-1 showed similar pathology to the mice infected with B.1.1.7 and this was also reflected by the pathology scores (**Figure 7A/B**). The lungs of mice infected with P.1 S in WA-1 showed increased inflammation compared to the lungs of mice infected with P.1 on both day 2 and day 4 (**Figure 7A/B**) postinfection. Despite B.1.351 S in WA-1 producing attenuated weight loss, the lung pathology scores for these mice were higher compared to mice infected with B.1.351, although none of the lung pathology differences were statistically significant (**Figure 7A/B**).

**Figure 7.**
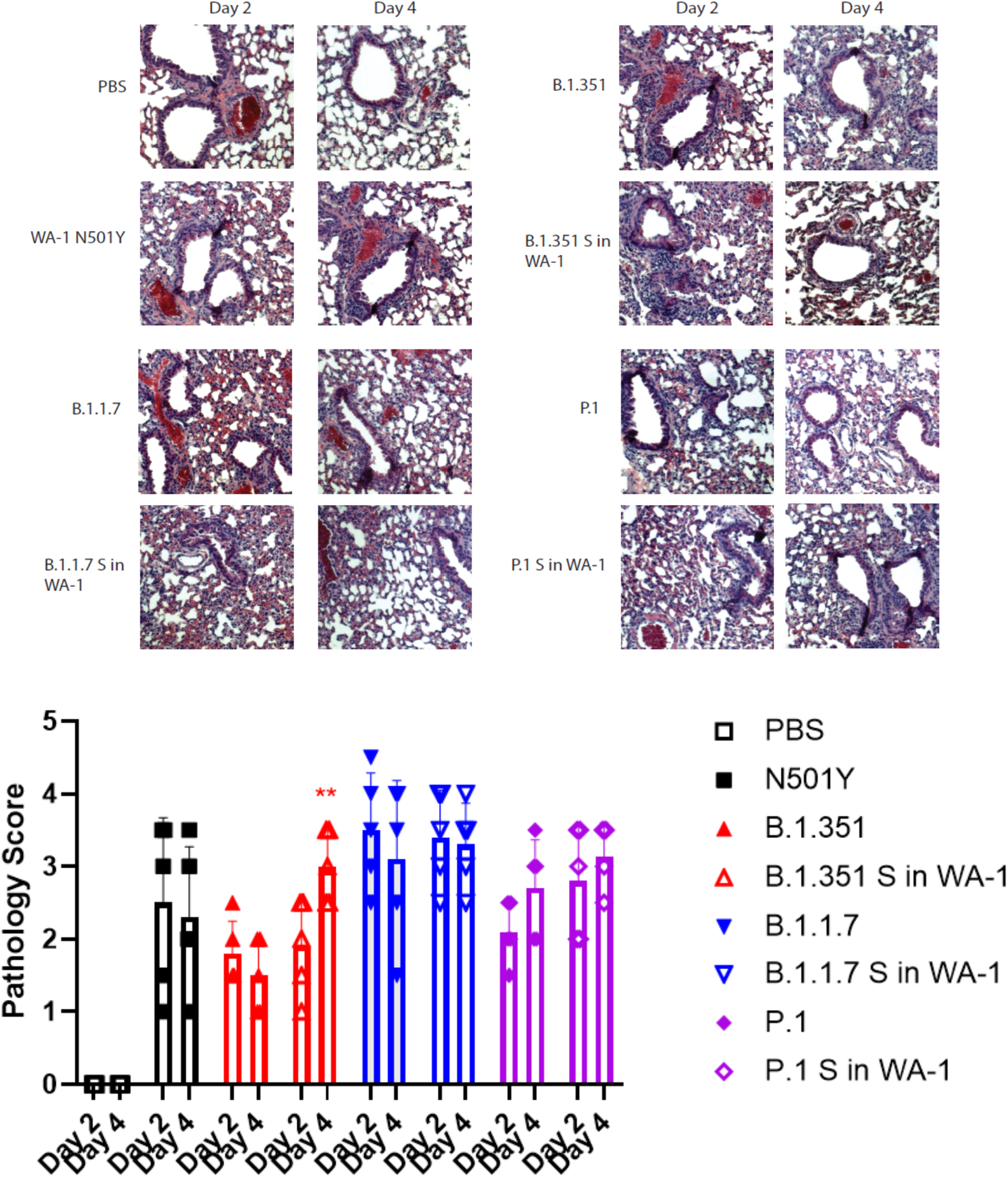
Lung pathology of 12-week old BALB/c mice infected with variant spike in WA-1 viruses and parent variants. A. H/E stained lung sections. B. Pathological scoring of the lungs.

### Lung Cytokine Profiling for WA-1, WA-1ΔORF3a, P.1, and P.1 S in WA-1

We then analyzed the host response to selected viruses that showed the most significant differences in weight loss and titer when compared to the parent strains. We compared the inflammatory cytokine and chemokine profile by qPCR on both the Day 2 lungs and the Day 4 lungs of the K18-hACE2 mice infected with WA-1, WA-1ΔORF3a/b, P.1, and P.1 S in WA-1 (**Table S7**). It is interesting to note that for both the WA-1ΔORF3a/b to WA-1 comparison and the P.1 S in WA-1 to P.1 comparison, the majority of significant cytokine and chemokine changes were seen at Day 2. For this reason, we will focus on the changes we saw at Day 2.

For K18-hACE2 mice infected with WA-1ΔORF3a/b, the majority of significant fold changes in cytokine and chemokine levels compared to mice infected with WA-1 were less than 1. The top five significantly downregulated genes are Ccl7, Csf3, Cxcl1, Cxcl3, and Cxcl5, which are key cytokines involved in neutrophil production and recruitment (**Table 1**). There was a small subset of genes which were significantly upregulated. These genes are Adipoq, Bmp7, Ctf1, Cxcl11, Il4, and Il5, which are mostly cytokines involved in cell survival and the early stages of T-cell activation (Table 1).

**Table 1.**
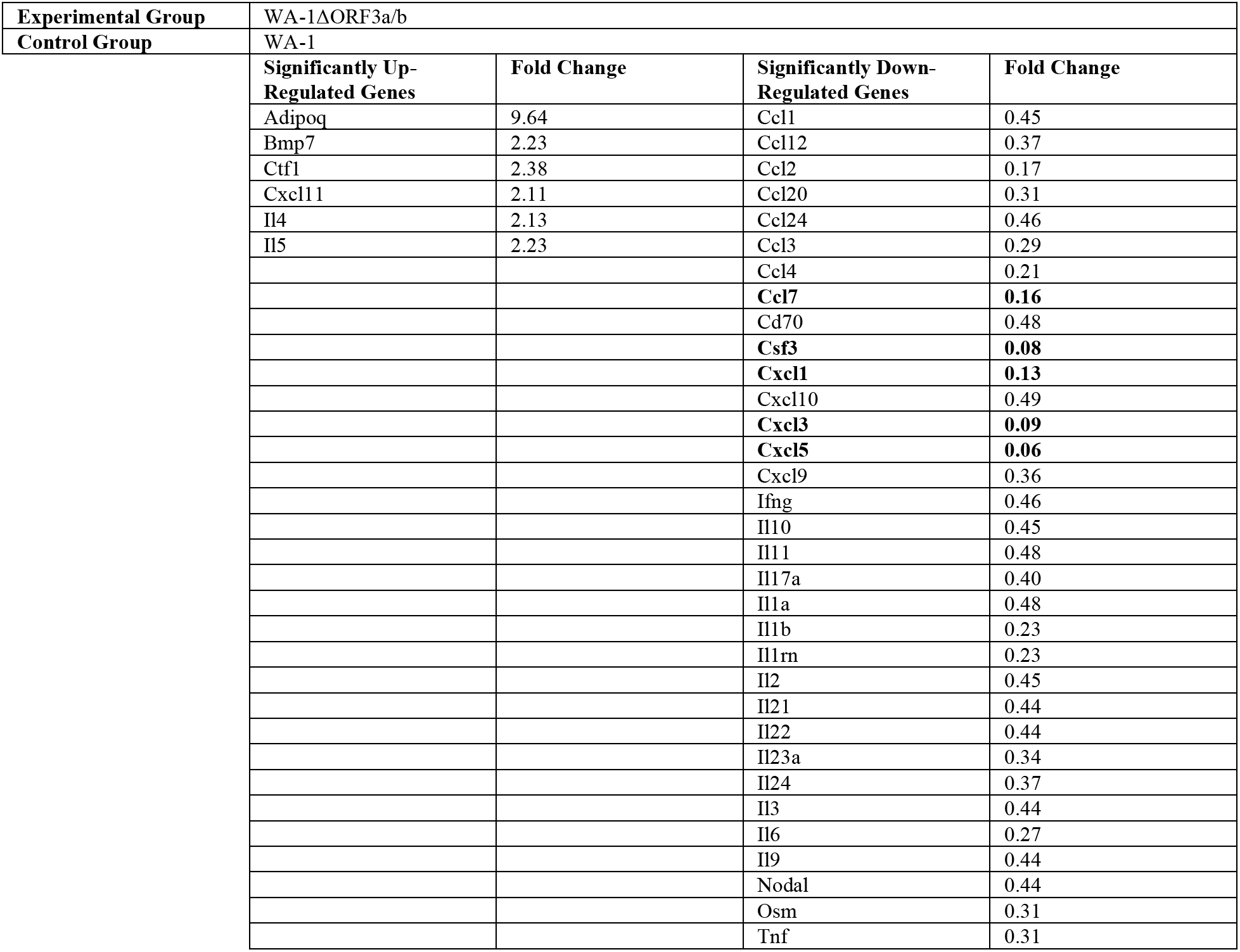
Fold changes of inflammatory chemokines and cytokines in Day 2 mouse lungs of mice infected with WA-1ΔORF3a/b compared to mice infected with WA-1.

As was seen with the WA-1ΔORF3a and WA-1 comparison, the majority of significant fold changes occurred at Day 2 for the P.1 and P.1 S in WA-1 comparison. There were few differences in gene induction when comparing P.1 and P.1 S in WA-1 in relation to the WA-1ΔORF3a/b and WA-1 differentially expressed genes. The top five upregulated genes were Ccl1, Ccl20, Csf3, Cxcl5, and Osm, which are involved in the attraction of neutrophils and lymphocytes (**Table 2**). All of the downregulated genes had similar fold changes. These genes include Csf2, Il12a, and Il21, which are known to be secreted by innate immune cells, including natural killer cells (**Table 2**).

**Table 2.** Fold changes of inflammatory chemokines and cytokines in Day 2 mouse lungs of mice infected with P.1 S in WA-1 compared to mice infected with P.1

| Experimental Group | P.1 S in WA-1 |  |  |  |
| --- | --- | --- | --- | --- |
| Control Group | P.1 |  |  |  |
|  | Significantly Up-Regulated Genes | Fold Change | Significantly Down-Regulated Genes | Fold Change |
|  | **Ccl1** | **2.91** | Adipoq | 0.36 |
|  | Ccl2 | 2.66 | Csf2 | 0.41 |
|  | **Ccl20** | **2.71** | Il12a | 0.35 |
|  | Ccl7 | 2.47 | Il21 | 0.46 |
|  | **Csf3** | **3.49** | Il5 | 0.48 |
|  | Cxcl1 | 2.47 | Mstn | 0.17 |
|  | **Cxcl5** | **4.23** | Thpo | 0.46 |
|  | Il1b | 2.02 | Tnfsf11 | 0.36 |
|  | Il1rn | 2.18 |  |  |
|  | Il23a | 2.65 |  |  |
|  | Il24 | 2.15 |  |  |
|  | Il6 | 2.47 |  |  |
|  | **Osm** | **2.72** |  |  |

## Discussion

A unique characteristic of coronaviruses is the inclusion of genes in the 3’ end of the genome that are unique to each coronavirus family.^16^ These genes are called “accessory genes” because, while unique to each family, they are not essential for replication.^17^ Many of these genes across the coronavirus family have been shown to alter host pathways including interferon signaling, cell cycle progression, and other assorted anti-viral responses to viral infection that enhance replication and pathogenesis.^9,10,18,19^

We designed two ways to analyze the functional consequences of the accessory genes in SARS-CoV-2 *in vivo*. First, we produced deletion viruses that deleted each ORF from our infectious clone of SARS-CoV-2. Second, we produced mutant SARS-CoV-2 viruses that contained the spike of each previously circulating variant. As variants of SARS-CoV-2 have emerged, the increasing incidence of mutations both within spike and outside of spike were noted. Although, the mutations in spike may enhance entry and kinetics of infection, the mutations observed in the other genes may alter disease severity through interactions with the host immune system. In our experiments we sought to differentiate between the role of spike mutations in each variant and those in other genes of the variant genome. As many of these nonspike mutations in the genome are in the accessory ORFs, we decided to couple our work with our variant spikes in WA-1 viruses with our work with accessory deletion viruses. This allows us to determine which accessory ORFs contribute significantly to pathogenesis through infection with our deletion viruses, and to link mutations in the corresponding variant ORFs to impacts on ORF function and pathogenesis.

Our work with the accessory deletion viruses has revealed that ORF3a and ORF3b contribute significantly to viral replication in K18-hACE2 mice. Mice infected with a WA-1ΔORF3a/b deletion virus demonstrated attenuated weight loss and significantly reduced lung titers at both day 2 and day 4. Our finding that an ORF3a/b deletion virus is attenuated in mice is supported by previously published work as does our finding that WA-1ΔORF6 is not attenuated in mice.^20^ This paper also found that deletions of ORF7a, ORF7b, and ORF8 show reduced lung titer, which we did not find in our experiments. This may be due to inoculation differences in our model where mice were infected with a non-lethal dose of 10^3^ pfu instead of 10^5^ pfu as used in the previous study. It is possible that we may see attenuation in weight loss and viral replication of WA-1ΔORF7a/7b, and WA-1ΔORF8 at higher doses or in different mouse backgrounds, which were not studied here.

The lung cytokine and chemokine profiles of mice infected with WA-1ΔORF3a/b demonstrated reduced expression of inflammatory cytokines and chemokines compared to WA-1. This difference is most likely attributed to the fact the WA-1ΔORF3a/b virus demonstrates attenuated replication in mice. As expected, the attenuated replication of WA-1ΔORF3a/b results in the downregulation of cytokines and chemokines involved in neutrophil recruitment, including Ccl7, Csf3, Cxcl1, Cxcl3, and Cxcl5. There is a small subset of genes that are upregulated in the WA-1ΔORF3a/b lungs. Two of these upregulated genes are Il4 and Il5, which function to drive the Th2 response. Interestingly, COVID-19 is known to skew to a Th2 response through stimulating the production of Il4 and Il5.^21^ Given that the WA-1ΔORF3a/b virus is attenuated compared to WA-1, we would expect a downregulation of these cytokines. The downregulation of Il4 might in part be explained by its role in the tissue remodeling process, with the faster clearance of WA-1ΔORF3a/3b allowing for tissue repair to take place.^22^ Another of these upregulated genes, adipoq, encodes the insulin-sensitizing hormone, adiponectin. Interestingly, reduced adiponectin levels are associated with severe respiratory failure in COVID-19 patients.^23^ Future work to characterize the role of these cytokines and chemokines SARS-CoV-2 pathogenesis will focus on how these pathways interact with the accessory genes of the virus.

As the emergence of variants of SARS-CoV-2 occurred, we noted the appearance of point mutations in the accessory genes of SARS-CoV-2. To elucidate the impact of non-spike mutations on viral pathogenesis, we synthesized variant spikes in the WA-1 backbone. We focused on the synthesis of the B.1.351, B.1.1.7, and P.1 lineages. When comparing the *in vitro* replication of the parent variants to their paired variant spikes in WA-1, we did not see many significant differences in replication in VeroE6 cells. The only significant difference we saw was a significant decrease in supernatant viral titer at 72 hours of P.1 S in WA-1 when compared to P.1.

In the K18-hACE2 mice, the absence of *in vitro* replicative differences for B.1.351 S in WA-1 and B.1.1.7 in WA-1 were reflected. Mice infected with B.1.351 in WA-1 exhibited no differences in weight loss, lung titer, and brain titer compared to mice infected with B.1.351. This was also true for mice infected with B.1.1.7 when compared to mice infected with B.1.1.7 S in WA-1. Results from the BALB/c experiment differed significantly for the B.1.351 S in WA-1/B.1.351 pairing, with B.1.351 S in WA-1 showing attenuated weight loss and lung titers compared to B.1.351. The mice infected with B.1.1.7 S in WA-1 showed an increase in lung titer on day 2, but no difference in weight loss was seen when compared to B.1.1.7.

The most significant replicative differences in both BALB/c and K18-hACE2 mice was seen with P.1 and P.1 S in WA-1. In both sets of mice, mice infected with P.1 S in WA-1 demonstrated increased weight loss and significantly higher lung titers by plaque assay and qPCR than mice infected with P.1. In the K18-hACE2 mice, the brain RNA and brain titers trended with this data as well, although the values were not significant.

Given the significant differences in P.1 S in WA-1 and P.1 replication *in vivo*, we analyzed cytokine and chemokine gene expression on both the day 2 and day 4 lungs of these mice. Compared to P.1, the P.1 S in WA-1 cytokine and chemokine profiled differed the most on Day 2. Most of the significant changes seen involved the upregulation of genes involved in neutrophil recruitment. The most upregulated gene was cxcl5, which has been implicated as the major chemoattractant for neutrophils in SARS-CoV-2 and a major cause of inflammation.^24^ It is therefore not unexpected that this gene would be upregulated due to the increased lung titer of P.1 S in WA-1 compared to P.1. The downregulated genes of interest include thpok and Il5, which are important for driving the differentiation of CD4^+^ T-cells. This finding could suggest that there is some inhibition of the CD4^+^ T-cell recruitment pathway by the non-spike genes in WA-1 that is attenuated or absent in P.1. There is also evidence of the loss of early interferon antagonism in the non-spike genes in P.1, as IFNγ levels in lungs from the P.1 S in WA-1 mice were reduced at day 2 compared to the P.1 mice despite the higher lung titers in the mice infected with P.1 S in WA-1. However, this difference was not seen at Day 4.

Together this work demonstrates that ORF3a/b have substantial roles in pathogenesis and host responses to SARS-CoV-2. It is interesting to note that two of the variant spike in WA-1 viruses studied here, B.1.351 and P.1, possess mutations in ORF3a. This points to the fact the emergence of mutations in the variant accessory ORFs, particularly ORF3a, contribute to viral pathogenesis. The possible advantageous effects on viral fitness and likelihood of transmission of these mutations is supported by the continued identification of mutations in these ORFs in new variants. We interpret this data to suggest that mutations outside of spike may be driving critical phenotypes of SARS-CoV-2 infection and disease. Although spike mutations may allow for better engagement of ACE2 to facilitate entry into cells as well as evasion of antibodies, the mutations in the accessory proteins may actually be negatively impacting disease leading to less severe clinical phenotypes. The balance of these two strategies may confer longer courses of virus replication and spread in the lungs while allowing for increased time for virus transmission from one infected person to their contacts. We hypothesize that this balance is critical for further evolution of SARS-CoV-2 and, as more variants emerge, we will identify additional mutations outside of spike that contribute significantly to viral replication, transmission, and pathogenesis.

## Acknowledgements

This work was supported by by grants from The Bill and Melinda Gates Foundation INV-006099, DARPA HR0011-20-2-0040, and DHS/BARDA ASPR-20-01495 to MF. Funding to SV was provided by NIH NIH R01AI137365, NIH R03AI146632 and the J Craig Venter Institute.

## Supplementary Figures

**Table S1.**
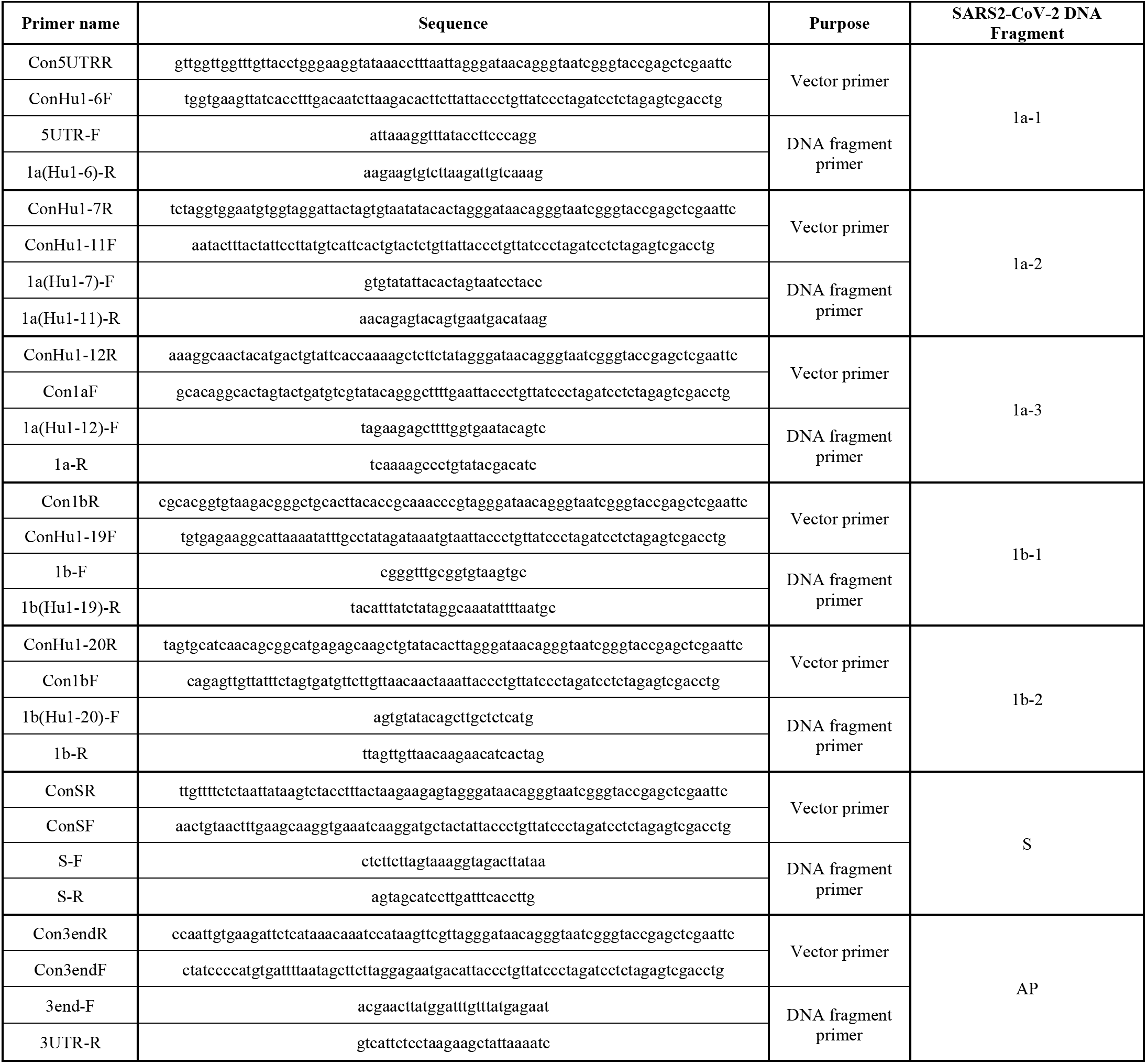
Primer sequences for cloning SARS-CoV-2 genome fragments.

**Table S2.**
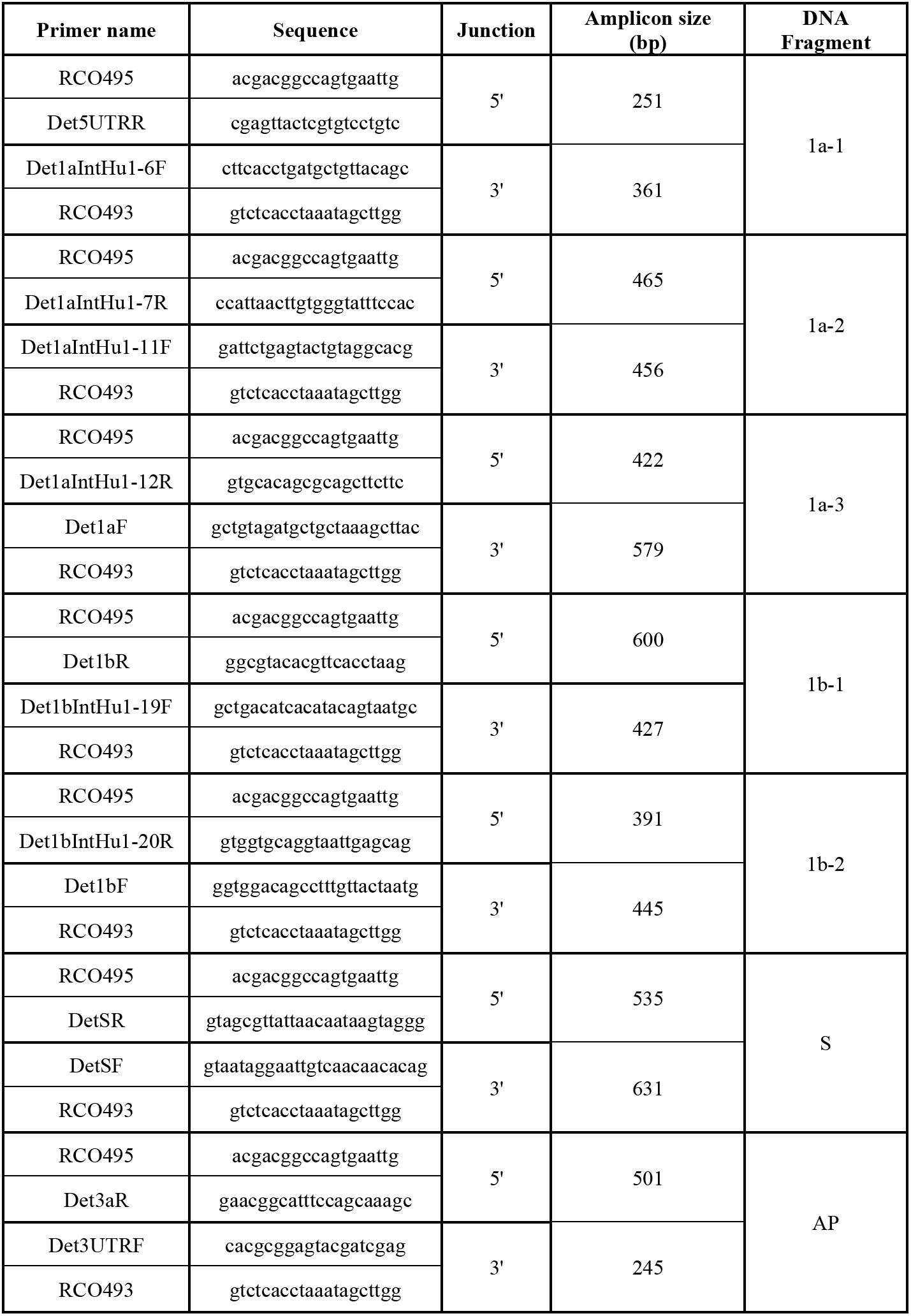
Detection primers to screen positive SARS-CoV-2 DNA fragment clones.

**Table S3.**
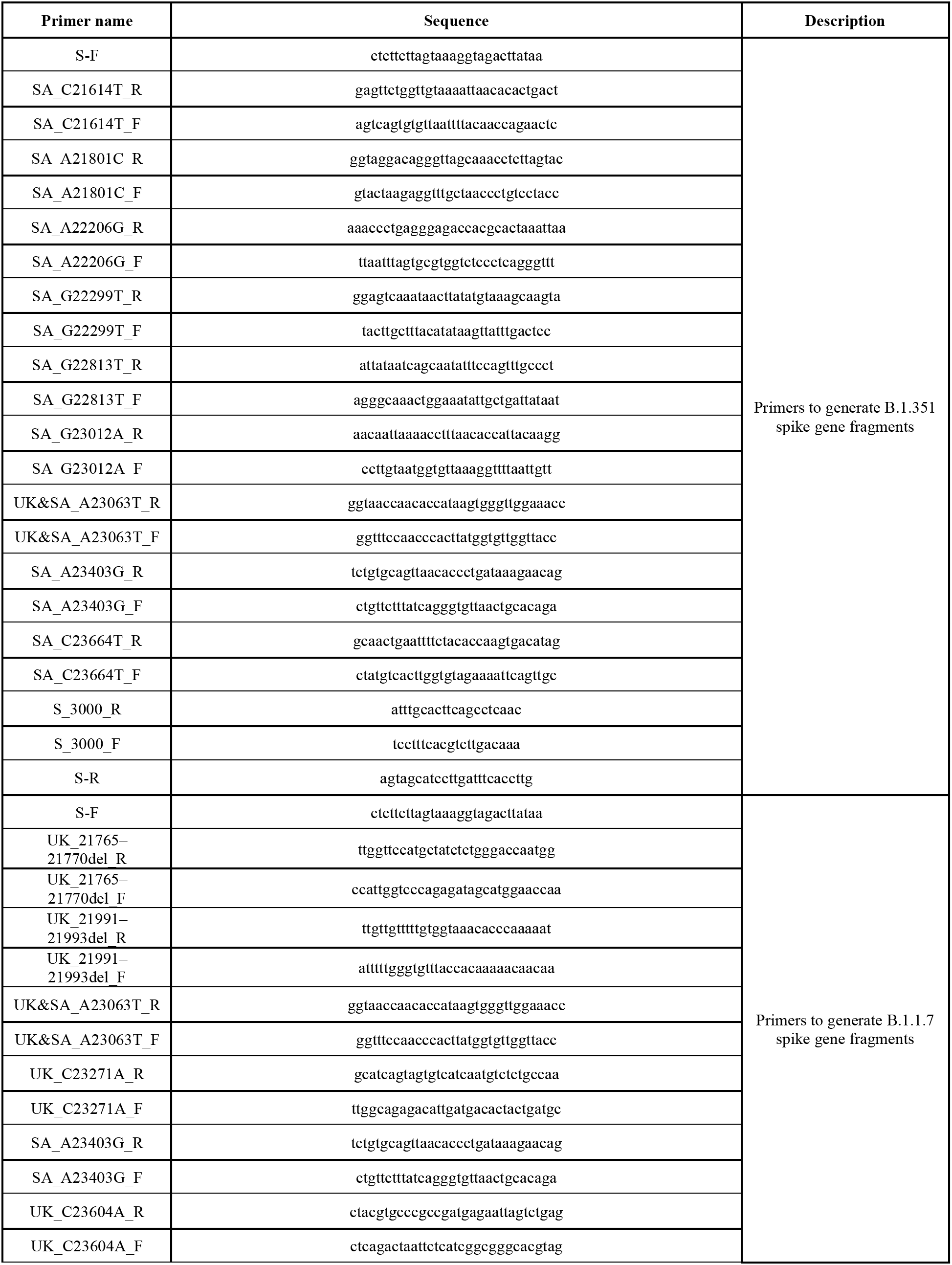
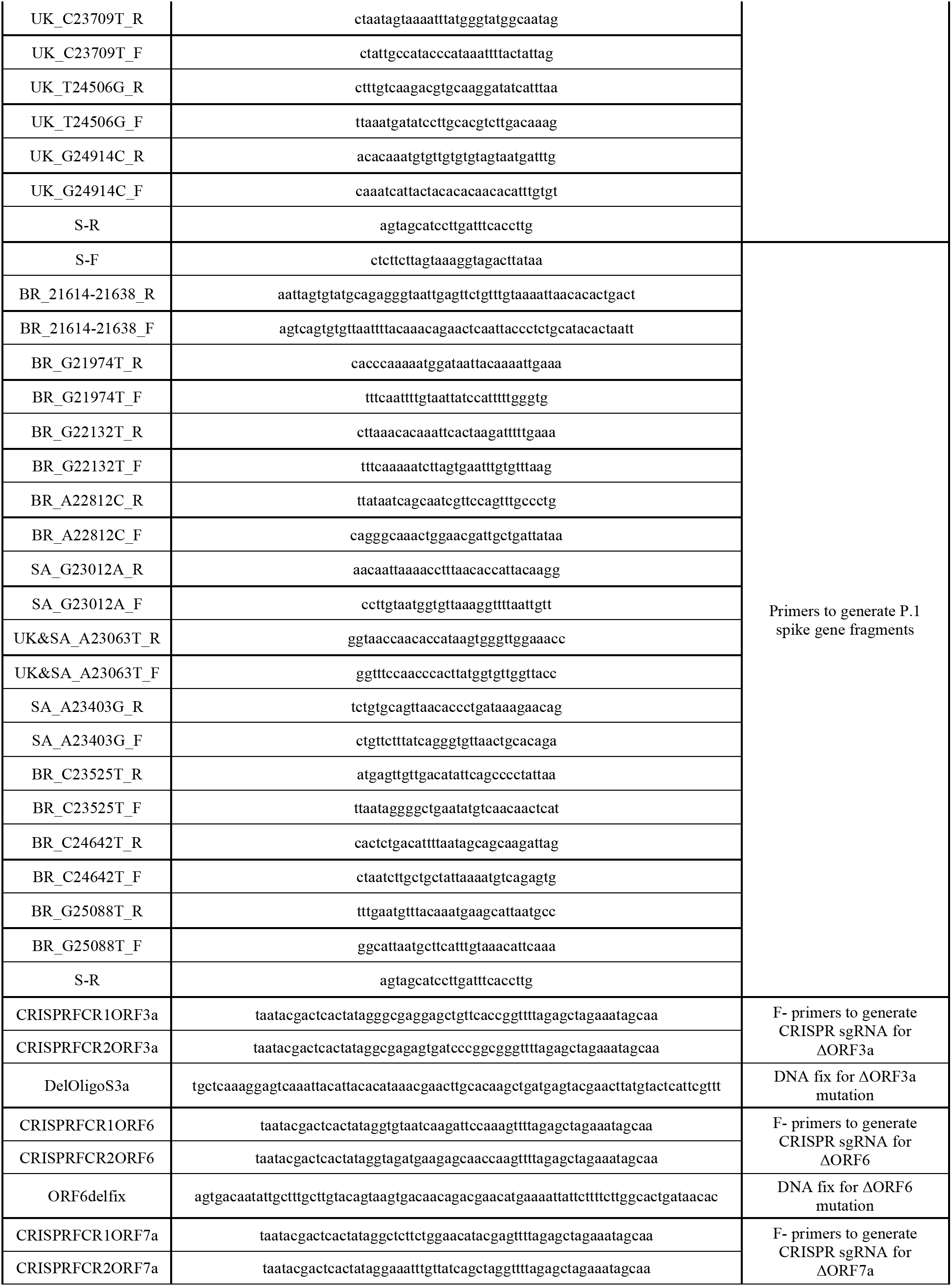
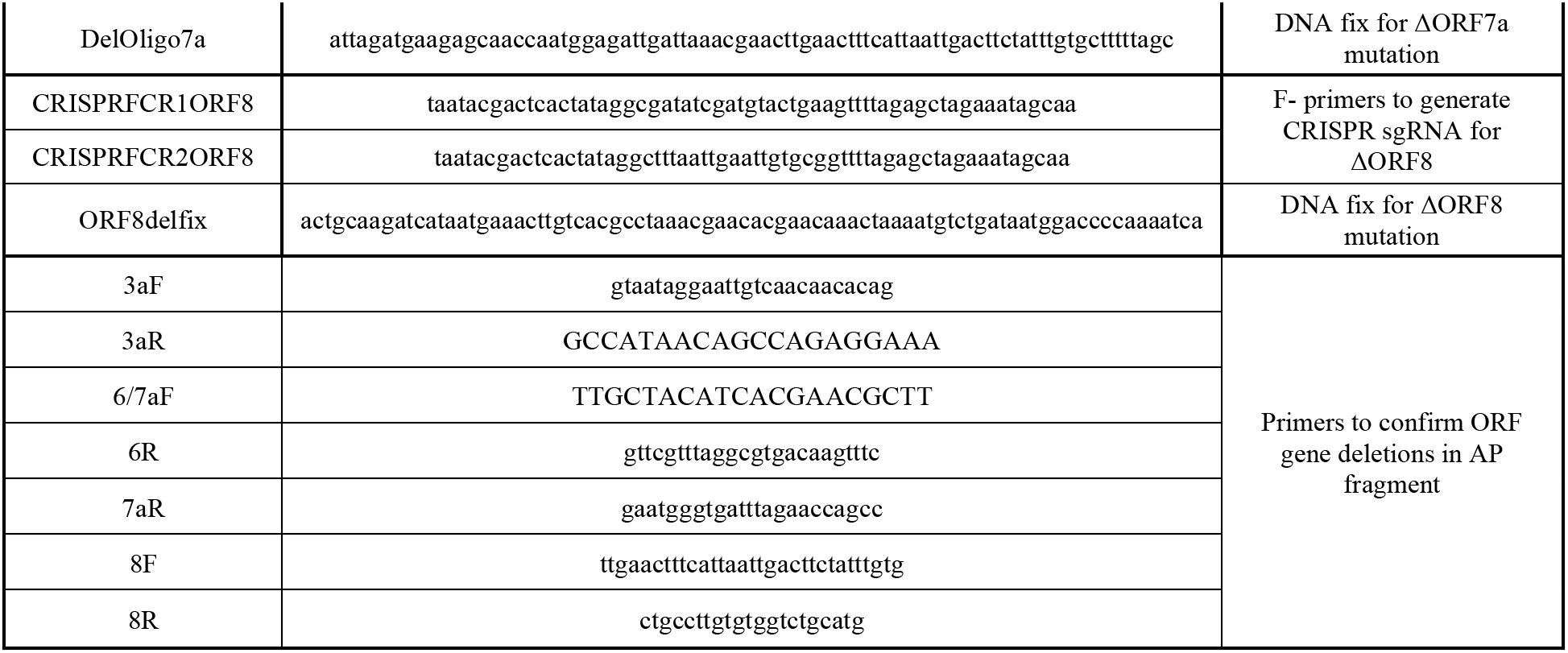
Primer sequences to generate and confirm SARS-CoV-2 mutants.

**Table S4.**
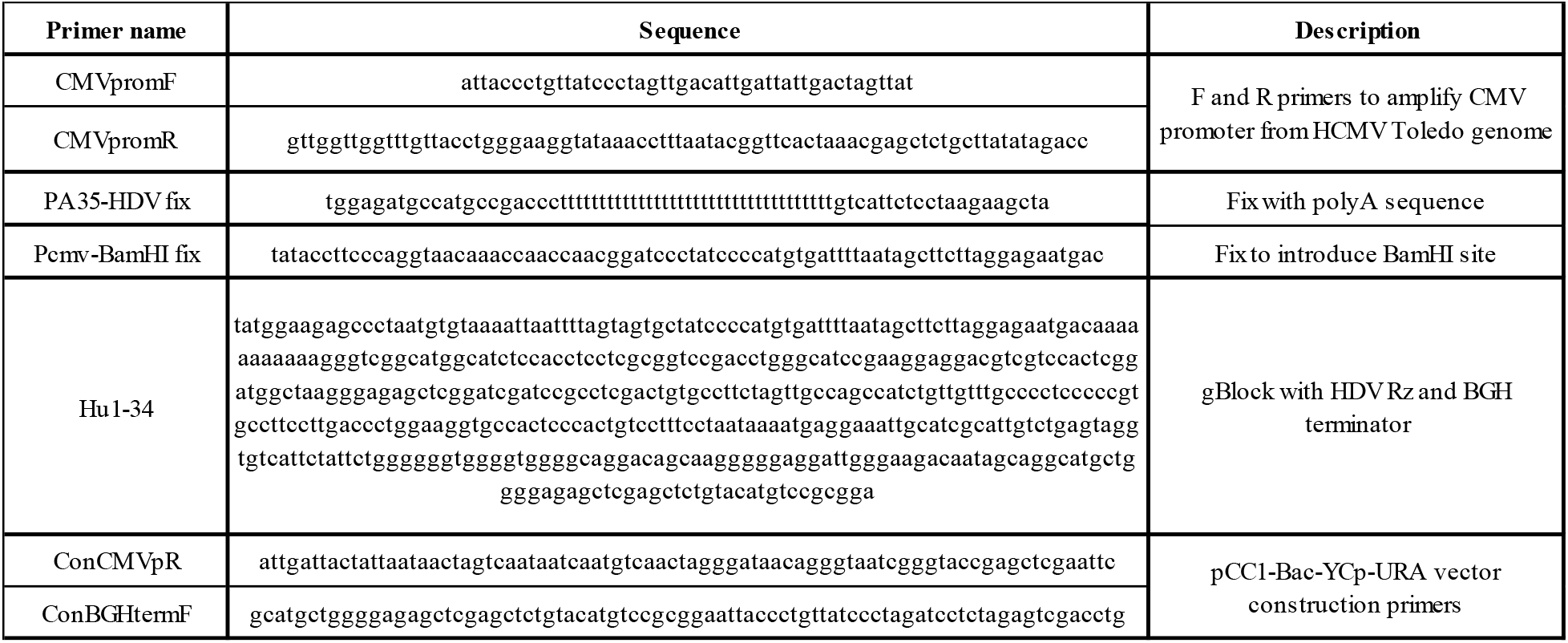
Primer sequences to construct SARS-CoV-2 complete genome vector.

**Table S5.**
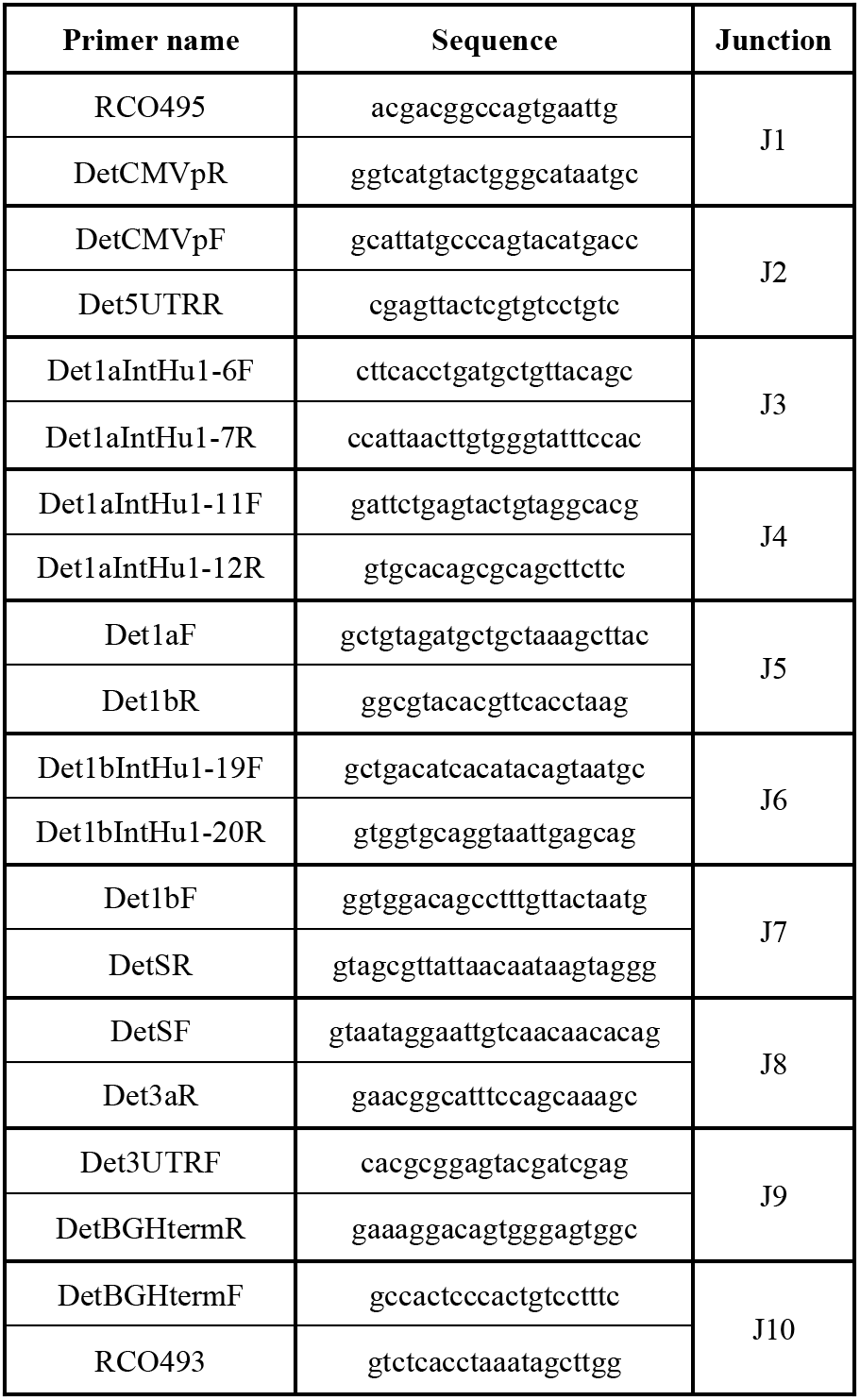
Detection primers to screen positive SARS-CoV-2 complete genomes.

**Table S6.** Variant spike mutations.

| Lineage | Variant spike amino acid substitutions vs WA1 |
| --- | --- |
| B.1.351 | L18F, D80A, D215G, R246I, K417N, E484K, N501Y, D614G, A701V |
| B.1.1.7 | HV 69–70del, Y144del, N501Y, A570D, D614G, P681H, T716I, S982A, D1118H |
| P.1 | L18F, T20N, P26S, D138Y, R190S, K417T, E484K, N501Y, D614G, H655Y, T1027I, V1176F |

**Table S7.**
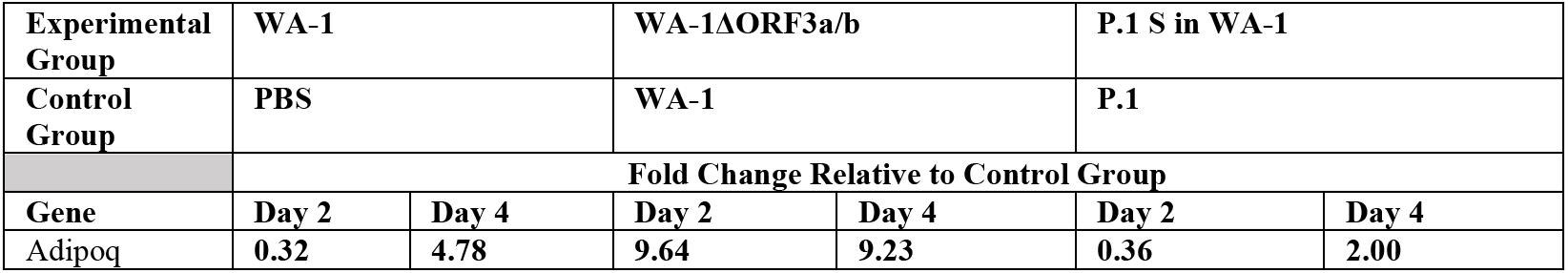
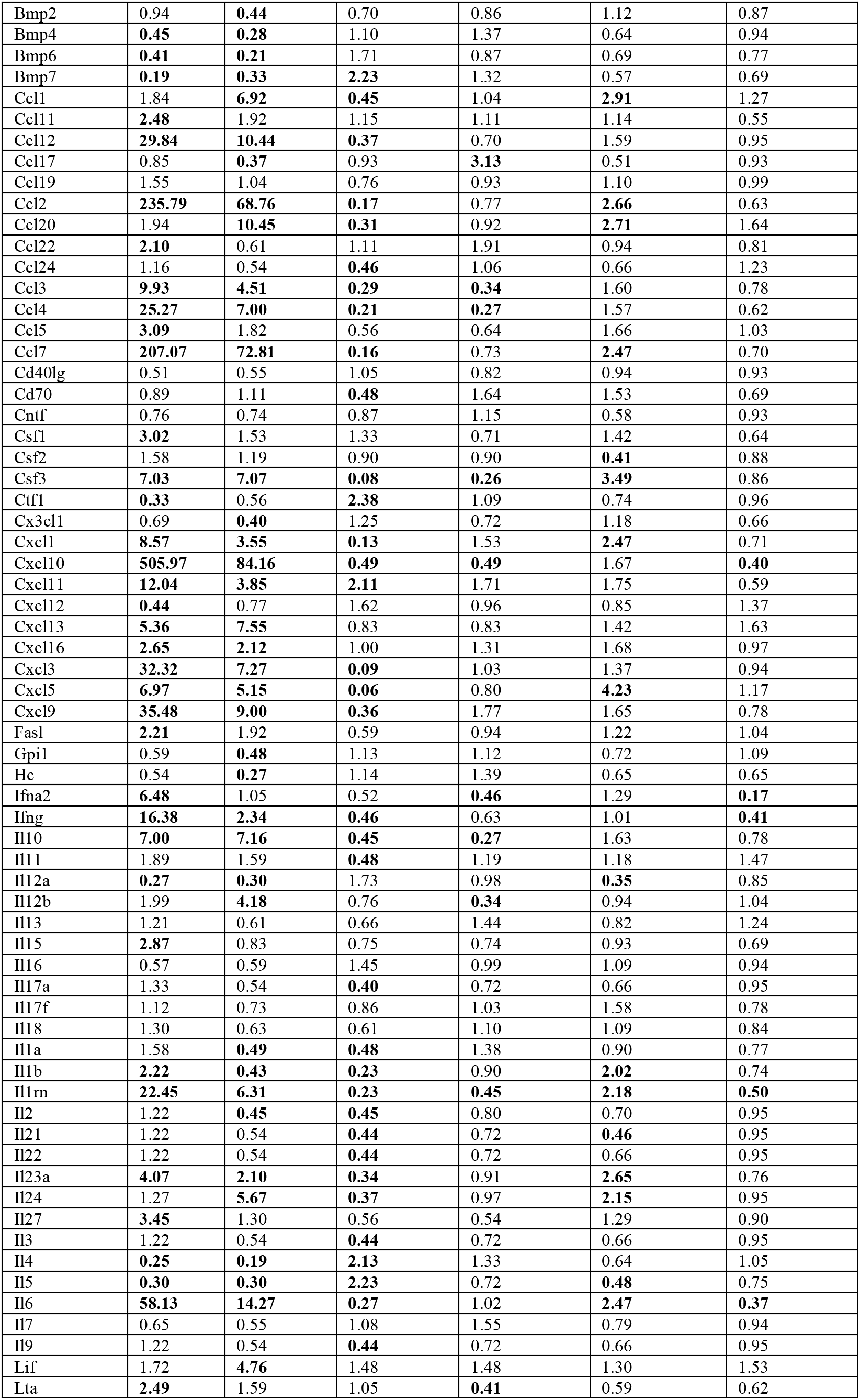
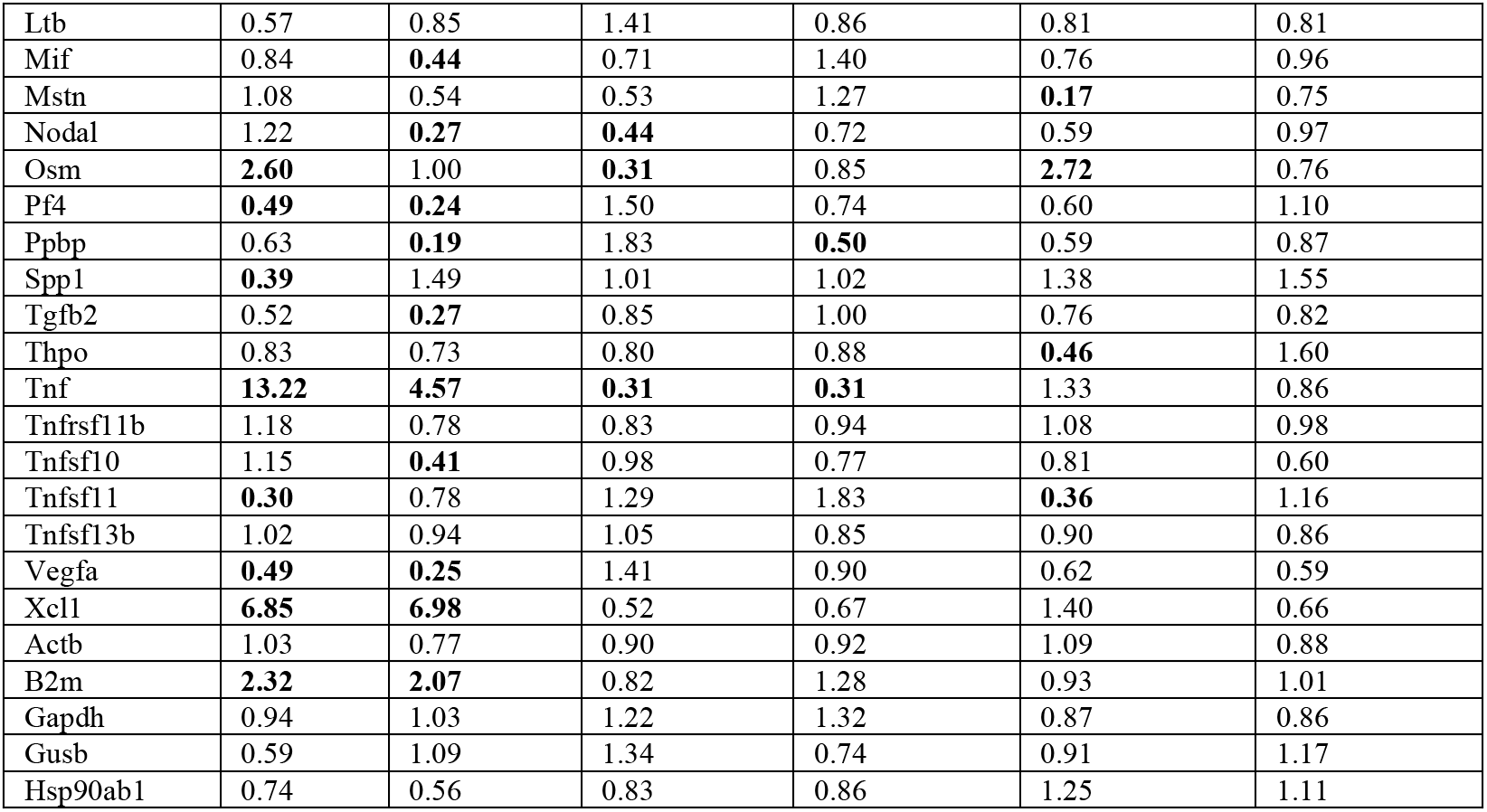
Fold changes of inflammatory cytokines and chemokines of mouse lungs on Day 2 and Day 4.

**Figure S1.**
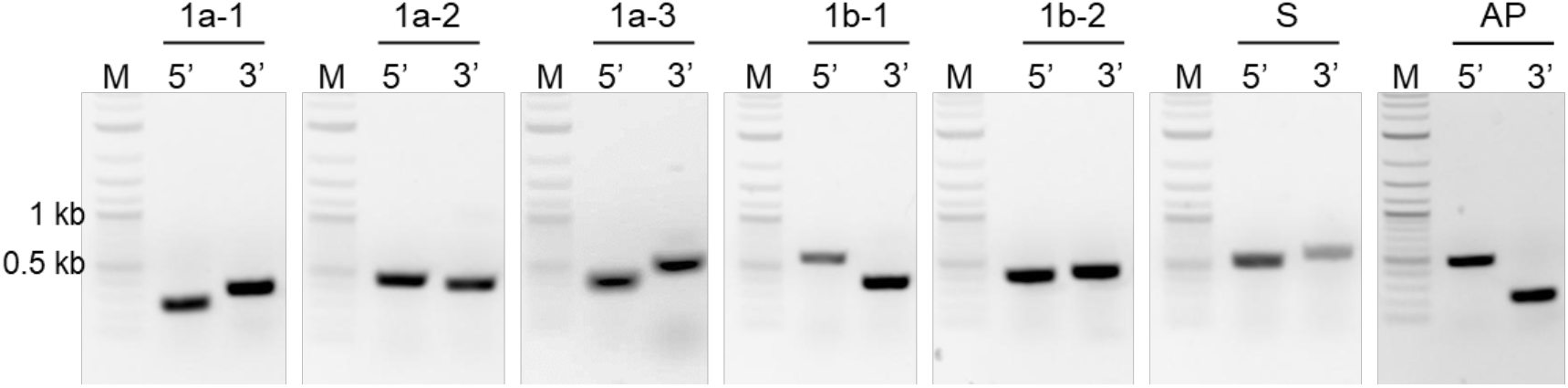
PCR confirmation of SARS-CoV-2 genome fragments used to assemble full-length genomes. Junctions between vector and each SARS-CoV-2 DNA fragment were PCR amplified using detection primers in Table S2. 5’, junction at 5’ end of SARS-CoV-2 fragment; 3’, junction at 3’ end of SARS-CoV-2 fragment; M: 2-log marker.

**Figure S2.**
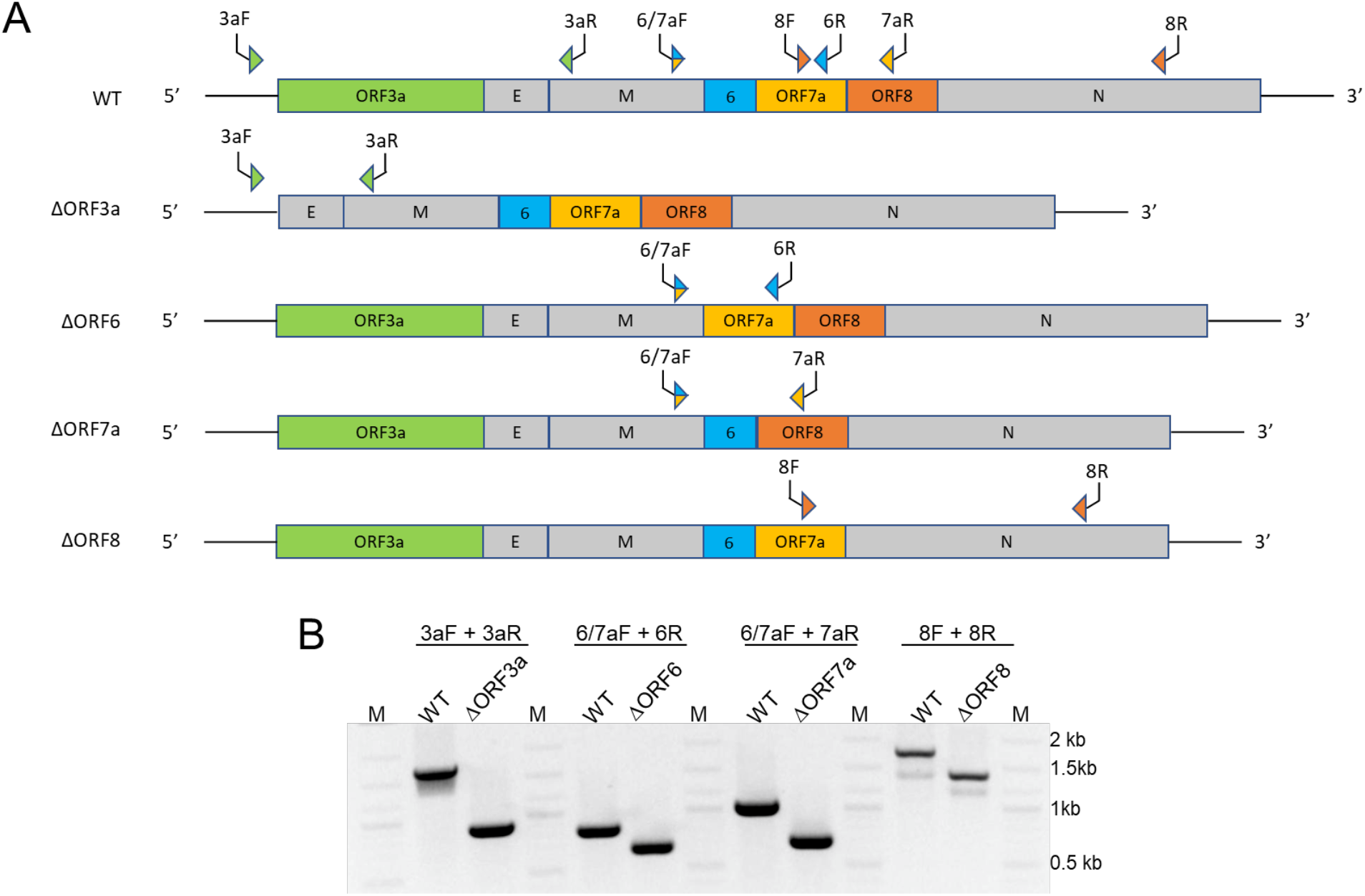
PCR amplification to confirm accessory ORF deletions. A. Schematic of SARS-CoV-2 WA1 accessory ORF deletion mutants (ΔORF3a, ΔORF6, ΔORF7a, ΔORF8). B. PCR screen to test for ORF deletions in mutants. Primer sequences are listed in Table S3. WT, wild type WA1; M: 2-log marker.

